# Tying the Knot: In Silico Design of Foldable Lasso Peptides

**DOI:** 10.1101/2025.01.17.633674

**Authors:** John D. M. Nguyen, Gabriel C. A. da Hora, Marcus C. Mifflin, Andrew G. Roberts, Jessica M. J. Swanson

## Abstract

Lasso peptides are a unique class of natural products with distinctively threaded structures, conferring exceptional stability against thermal and proteolytic degradation. Despite their promising biotechnological and pharmaceutical applications, reported attempts to prepare them by chemical synthesis result in forming the nonthreaded branched-cyclic isomer, rather than the desired lassoed structure. This is likely due to the entropic challenge of folding a short, threaded motif prior to chemically mediated cyclization. Accordingly, this study aims to better understand and enhance the relative stability of pre-lasso conformations—the essential precursor to lasso peptide formation—through sequence optimization, chemical modification, and disulfide incorporation. Using Rosetta fixed backbone design, optimal sequences for several class II lasso peptides are identified. Enhanced sampling with well-tempered metadynamics confirmed that designed sequences derived from the lasso structures of rubrivinodin and microcin J25 exhibit a notable improvement in pre-lasso stability relative to the competing nonthreaded conformations. Chemical modifications to the isopeptide bond-forming residues of microcin J25 further increase the probability of pre-lasso formation, highlighting the beneficial role of non-canonical amino acid residues. Counterintuitively, the introduction of a disulfide cross-link decreased pre-lasso stability. Although cross-linking inherently constrains the peptide structure, decreasing the entropic dominance of unfolded phase space, it hinders the requisite wrapping of the N-terminal end around the tail to adopt the pre-lasso conformation. However, combining chemical modifications with the disulfide cross-link results in further pre-lasso stabilization, indicating that the ring modifications counteract the constraints and provide a cooperative benefit with cross-linking. These findings lay the groundwork for further design efforts to enable synthetic access to the lasso peptide scaffold.

**SIGNIFICANCE:** Lasso peptides are a unique class of ribosomally synthesized and post-translationally modified natural products with diverse biological activities and potential for therapeutic applications. Although direct synthesis would facilitate therapeutic design, it has not yet been possible to fold these short sequences to their threaded architecture without the help of biosynthetic enzyme stabilization. Our work explores strategies to enhance the stability of the pre-lasso structure, the essential precursor to *de novo* lasso peptide formation. We find that sequence design, incorporating non-canonical amino acid residues, and design-guided cross-linking can augment stability to increase the likelihood of lasso motif accessibility. This work presents several strategies for the continued design of foldable lasso peptides.

## INTRODUCTION

Lasso peptides are a fascinating class of natural products with unique structural features. They are characterized by a threaded, knot-like [1]rotaxane arrangement, defined by a macrocyclic ring that encircles a C-terminal peptide tail to produce distinct loop, ring, and tail components (**Figure 1A**). Their threaded structure imparts exceptional stability, making lasso peptides highly stable against chemical, thermal, and proteolytic degradation. This stability, coupled with their relatively small, compact structure and diverse biological activities—such as antimicrobial, anticancer, and enzyme inhibitory properties (1-4)—that positions lasso peptides as promising candidates for biotechnological and pharmaceutical applications.

**Figure 1.**
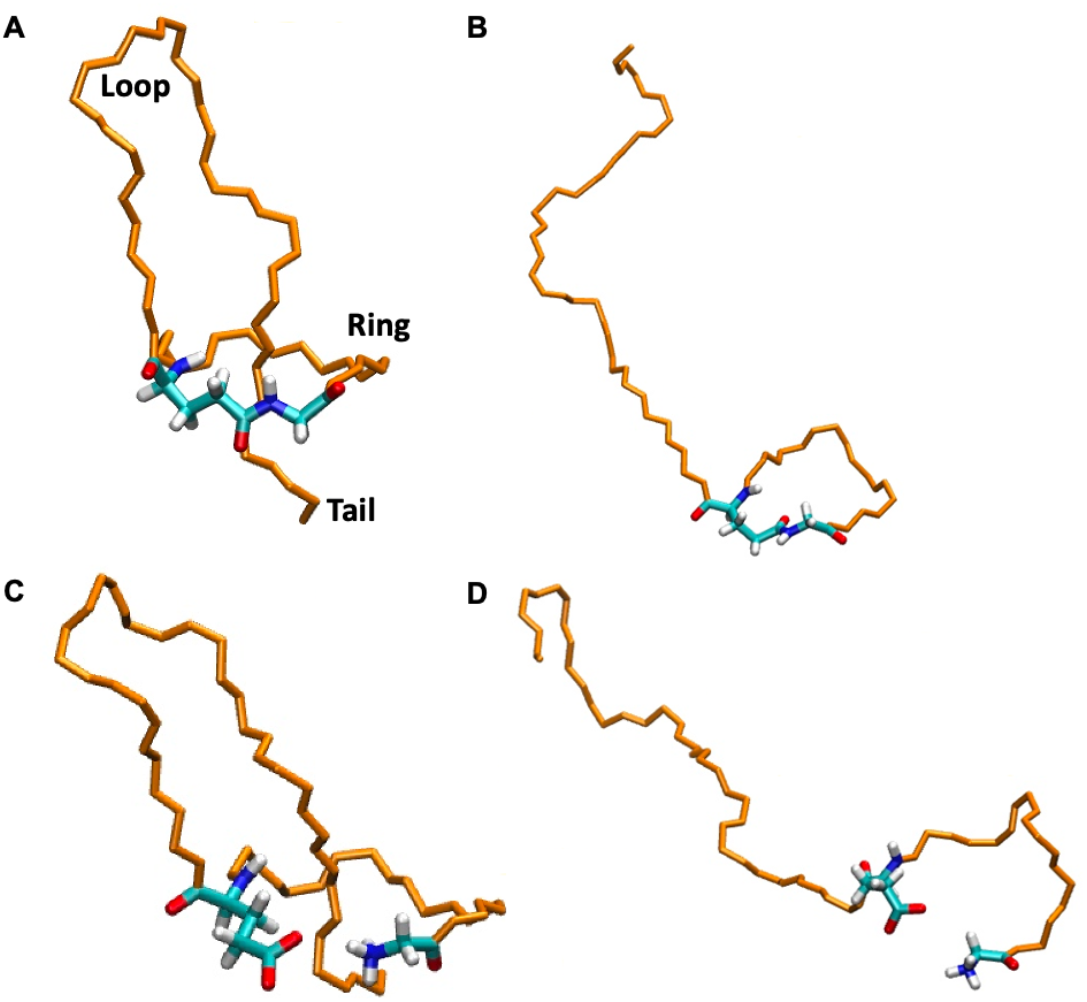
Upon formation of the isopeptide bond, lasso peptides get trapped in either the lasso (A) or tadpole (B, nonthreaded) conformation. Before the isopeptide bond forms, the pre-ring conformations are referred to as pre-lasso (C) and pre-tadpole (D). The essential isopeptide bond-forming residues are colored by atom type, and the remaining peptide residues are colored orange.

While the class II threaded motif confers stability and biologically important activities, achieving this specific arrangement by chemical synthesis through cyclization of an acyclic precursor remains challenging and has, to date, only led to the formation of the nonthreaded isomer, referred to herein as the tadpole conformation (**Figure 1B**) (5-9). Natural lasso peptides are formed by a lasso cyclase enzyme-dependent step that uses ATP to activate a sidechain carboxylate followed by irreversible cyclization upon the formation of an isopeptide bond between the N-terminal amine and side chain carboxylic acid of residue 7, 8, or 9 (**Figure 1A**) (5,10-12). Prior to cyclization, the native peptide must first be arranged into an enzyme-stabilized pre-lasso conformation (**Figure 1C**). Recent work predicted key interactions between the adenylated intermediate and its cyclase that can be altered to increase substrate tolerance in cell-free biosynthesis of lasso peptides (13). Indeed, the development of various biochemical approaches has greatly expanded the scope of accessible lasso peptides, highlighting the impressive range of natural substrate tolerance that has been characterized for lasso cyclases toward enabling the future design of functional lasso peptides (14-28). The information gained through bioinformatics tools such as RODEO (29) and high-throughput screening (16,21,22) of lasso peptide variant production is enabling the development of new tools for the prediction of lasso cyclase substrate compatibility and improvement of lasso peptide biological activity (30), as well as strategies for engineering lasso cyclase enzymes for increasing the tolerated substrate variance (13). Collectively, these results suggest that association with the lasso cyclase plays an essential role in stabilizing the pre-lasso conformation in natural sequences since cyclization occurring prior to the adoption of a pre-lasso conformation would yield the tadpole isomer. While there are exceptions (31), in most studied cases the lasso motif provides superior biological function and potency (32-35).

The elusive nature of a synthetic lasso motif has prompted computational investigations into the intrinsic (or *de novo*) folding of lasso peptides (36-38). Computational methods have become valuable tools for exploring various aspects of lasso peptides, including structure prediction (39,40), conformational stability (41), receptor binding (42), thermal unthreading, (43) cyclase design (13), and substrate prediction (44). Seminal computational work by Ferguson et al. (6) studied the folding propensity of the lasso peptide microcin J25 (MccJ25) (7) and found that it more readily adopted a left-handed pre-lasso conformation, in which the N-terminus wraps around the tail in a counterclockwise direction (45). This was an unexpected finding as natural MccJ25 is right-handed (7) (**Figure 1A,C**). The folding behavior of MccJ25 was later revisited with updated forcefields, revealing that the peptide indeed adopts the natural right-handed pre-lasso conformation (37). Despite expert sampling and analysis, the previous observation of the left-handed pre-lasso structure was attributed to the use of the GROMOS96 43A2 forcefield (46), which has since been updated to correct backbone dihedral parameters (47-50).

While simulations of MccJ25 demonstrated the formation of the right-handed pre-lasso fold was possible (37), they also revealed that it is rare—with the peptide readily unraveling to the entropically favored unfolded and pre-tadpole conformations (**Figure 1D**). Recent work identified collective variables (CVs) that can be used to characterize and quantify the folding propensity of MccJ25 via enhanced free energy sampling (38). The resulting free energy profiles and pre-lasso vs. pre-tadpole probabilities highlighted a substantial energetic preference for the pre-tadpole ensemble over the pre-lasso. This observation suggests that *de novo* pre-lasso folding will likely require increased stabilization of the pre-lasso fold. Encouragingly, there have been findings that hint at the feasibility of achieving such stabilization. By incorporating thiol auxiliary (at residue 1) and thioester (at residue 8) functionalities onto the isopeptide bond-forming residues of MccJ25, a significant increase in the percentage of pre-lasso conformations is observed in standard molecular dynamics (MD) simulations (37). Furthermore, it has been shown that protonation of the His5 imidazole-bearing side chain of the residue-modified MccJ25 provided additional stabilization to the pre-lasso ensemble (37).

Here, we build upon the gained insights and established tools to explore strategies for enhancing pre-lasso stability, ultimately aiming to facilitate synthetic access to the lasso motif. Our previous findings suggested that the pre-lasso fold can be stabilized through sequence optimization, unnatural modifications, and specific reaction conditions, such as pH (37). In this study, we employ sequence optimization through fixed backbone design using Rosetta (51) for a subset of class II lasso peptides. Class II lasso peptides are particularly interesting due to their lack of disulfide bonds that would lower the entropic freedom of the unfolded state, wherein the isopeptide forming residues are too far apart to cyclize and the unrestricted loop/tail regions can readily dissociate prior to encirclement by the pre-ring. However, class II lasso peptides may possess greater inherent stability to overcome the unfolded entropy and are the most prevalent class formed in nature. The best scoring designed sequences, along with their native counterparts, are compared using Well-Tempered Metadynamics (WTMetaD) with previously identified CVs (38) to quantify the efficacy of sequence optimization. The impact of sequence design is also combined with chemical modifications, showing that stabilization of the isopeptide-forming residues through non-canonical modifications has an additive effect on pre-lasso stability. Finally, we test the value of reducing the vast unfolded conformational phase space of lasso peptides by introducing a cross-link in the form of a disulfide bond into the designed sequences. The observed stabilization effects demonstrate that access to the pre-lasso conformation can be markedly improved and provides guidance for future efforts to design lasso peptides that are accessible through chemical synthesis and *de novo* folding.

## METHODS

### Rosetta

Rosetta is a versatile suite of algorithms for macromolecular modeling, including protein structure prediction, design, and docking (51). Its design capabilities have been instrumental in optimizing the stability and function of protein structures (52-55). RosettaScripts (56) was used for local minimization and sequence design of class II lasso peptides. Four design strategies were first employed with native MccJ25 to identify an optimal protocol: 1) fixed-backbone design (`PackRotamersMover`) with lasso structures, 2) `PackRotamersMover` with pre-lasso structures, 3) Flexible-backbone design with alternating rounds of design and backbone minimization (`FastDesign`) with lasso structures, and 4) `FastDesign` with pre-lasso structures. Of the four strategies, sequences generated by strategy 1 maintained the pre-lasso state the longest before fully unfolding in standard MD simulations. Consequently, strategy 1 was selected for application to the remaining peptides. Prior to design, a local energy minimization was performed with each NMR solution or crystal structure using `FastRelax`. REF2015 was used as the score function with distance, angle, and dihedral constraints to maintain the lasso isopeptide bond for strategies 1 and 2. Design was performed using the same score function and constraints as minimization. `ReadResfile` was used to control sampling at each residue position. One thousand design trajectories were run for each lasso structure, a number that was found to consistently give the best scoring sequence for each structure. `Disulfidize` was used to identify and build a disulfide into the native MccJ25 sequence. A sample RosettaScripts routine used for fixed-backbone design is shown in **Listing S1**.

### Well-Tempered Metadynamics

Metadynamics (MetaD) (57,58), a well-established enhanced free energy sampling method, was used to quantify the relative probability of each sequence reaching the pre-lasso vs. pre-tadpole ensembles. MetaD enables the investigation of physical processes occurring over time scales that are typically inaccessible with conventional MD simulations. MetaD achieves this by applying a history-dependent bias, usually in the form of a Gaussian function, along predefined CVs. This bias is periodically introduced into the system, modifying the Hamiltonian in such a way that the system is encouraged to move out of energy wells, thereby exploring new regions of the phase space.

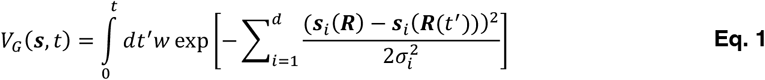

The Gaussian bias in MetaD is characterized by several parameters (Eq. 1): the energy rate *w*, which is determined by the height of the Gaussian (*w*_0_) divided by the frequency of deposition (τ); the Gaussian width σ_*i*_ for the *i*-th CV; and ***s***_#_(***R***(*t*^″^)), the value of the *i*-th CV at time *t′*. By integrating the negative of the applied MetaD bias, the underlying free energy surface along the chosen CVs can be reconstructed. However, a limitation of traditional (non-tempered) MetaD is the constant height of the Gaussians throughout the simulation, which can cause the free energy estimates to oscillate around their true values and potentially destabilize the system due to excessive energy input. To mitigate this issue, WTMetaD (59) introduces a time-dependent bias potential, where the height of the Gaussian decreases exponentially as a function of the local bias energy (Eq. 2). This adjustment is governed by a parameter Δ*T*, which controls the rate of decrease in Gaussian height. This method ensures that the bias potential converges asymptotically to a linear scaling inverse of the free energy (60), thus providing a more accurate and stable sampling of the system’s energy landscape.

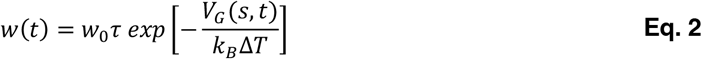

### Harmonic linear discriminant analysis (HLDA)

Harmonic linear discriminant analysis (HLDA) is an essential component in the previously identified protocol for defining lasso peptide CVs (38). Specifically, HLDA identifies a decision boundary that maximally separates the pre-lasso region of phase space from both pre-tadpole and unfolded conformations. HLDA builds upon linear discriminant analysis (LDA) (61), which is a supervised dimensionality reduction method that projects input data onto a new subspace, maximizing the Fisher objective function ***J*** to find a decision boundary with maximal class separability. The objective function (Eq. 3)

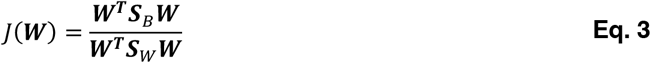

involves the within-class scatter matrix ***S***_*W*_, the between-class scatter matrix ***S***_*B*_, and the transformation matrix ***W*** that defines the subspace. Maximizing ***J*** is achieved by solving the generalized eigenvalue problem 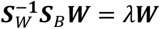. Despite its widespread use in classification, classical LDA has a significant limitation in that each pair of classes contributes equally to the between-class variance via the arithmetic mean, which causes larger between-class distances to dominate (6). This approach can be counterintuitive from a data analysis perspective, as classes with smaller variances should be better defined. From a rare-event perspective, it might be more appropriate to give greater weight to states with smaller fluctuations. To address this issue, a different measure for the within-class scatter matrix using the harmonic average was proposed (Eq. 4) (62,63). For a system with two metastable states, A and B, the collective variables *s* is expressed as:

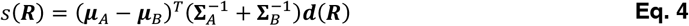

where ***μ***_*i*_ is the mean vector of class *i*, **Σ**_*i*_ is the class scatter matrix for metastable state *i*, ***d*** is the descriptor vector, and ***R*** represents the atomic coordinates. In HLDA metadynamics, the system is encouraged to transition from metastable state A to state B in a direction perpendicular to the decision boundary ***W*** that separates these states.

### Simulation Protocol

Molecular dynamics (MD) and MD enhanced via WTMetaD simulations of the lasso peptides rubrivinodin, caulosegnin II, achromonodin-1, and MccJ25, and their designed sequences were performed using the all-atom AMBER ff19SB force field (64). Initial configurations were solvated in periodic octahedron boxes using the OPC water model (65). A minimum 10 Å distance between the solute and the box edge was maintained, and each system was neutralized with Na^+^ or Cl^-^ ions if needed. The systems underwent minimization via the steepest descent method, initially with heavy atom restraints on the peptide (10 kcal mol^-1^ Å^2^ harmonic positional restraint for 1000 steps), which were gradually released without using the SHAKE algorithm (66,67). Equilibration involved eight steps, progressively reducing backbone atom restraints. The SHAKE algorithm (66,67) was introduced in the second step to constrain hydrogens. The first four steps employed the NVT ensemble (constant number of particles, volume, and temperature), while the last four used the NPT ensemble (constant number of particles, pressure, and temperature). The first five steps were each 5 ps with a 1 fs time step. The final three stages, each lasting 10 ps with a 2 fs time step, decreased the backbone restraint force constant from 1.0 to 0.5 kcal mol^−1^ Å^−2^, eventually removing positional restraints. Temperature was maintained at 300 K using the weak-coupling algorithm, and isotropic pressure scaling was applied for the NPT simulations, maintaining a constant pressure with a relaxation time of 1 ps.

In the production phase, each sequence was first simulated in standard MD to obtain structural ensembles for the HLDA analysis. A 2-fs time step was used, and simulations were run for 2 μs. The Langevin thermostat (68) maintained system temperature at 300 K, with a collision frequency of 1 ps^−1^. Pressure was kept at 1 bar using the Berendsen barostat (69), with a relaxation time of 1 ps. Long-range electrostatic interactions were calculated using the particle mesh Ewald (PME) method with an 8 Å nonbonded cutoff. Van der Waals interactions were treated with an 8 Å cutoff, consistent with the PME real space cutoff. Coordinates and energies were recorded every 5000 steps. Following standard MD, a triangulation-based algorithm, Lasso Threading Classification and Handedness Evaluation Descriptor (LATCHED) (37), was used to identify pre-lasso and pre-tadpole conformations by defining triangles between consecutive C-alpha atoms on residues 1−8 and determining if the tail-piercing vectors fell within any triangles. The LATCHED code is available on GitHub at https://github.com/gabedahora/LATCHED. Pre-lasso, pre-tadpole, and unfolded structures obtained from standard MD were then included in the HLDA analysis to define a CV for each sequence.

In the subsequent biased simulations, three replicas were simulated for each sequence, using the same parameters as standard MD simulations and averaging ∼1200 ns of simulation time per replica—totaling approximately 33.6 μs. Biased simulations continued until the bias height became negligible. All WTMetaD simulations were carried out using Amber 2020 (70) patched with PLUMED 2.7.1 (71,72). A Gaussian hill height of 0.03 with a bias factor of 10 was used to ensure efficient sampling of the pre-lasso, pre-tadpole, and unfolded regions of phase space while also maintaining the resolution of the energy basins. The width of the Gaussian hills varied for each sequence as the value corresponded to half of the standard deviation of each CV in standard MD. The free energy landscape of each replica was assessed at 500 ns intervals, and convergence was considered achieved once no significant changes were observed in the landscape. A sample PLUMED input file used for WTMetaD is shown in **Listing S2**. Analysis was conducted using CPPTRAJ (73), and 2D free energy surfaces were generated with custom Python scripts. Structure visualization and image generation were done using VMD 1.9.3 (74).

## RESULTS AND DISCUSSION

### Sequence design

An outstanding question that motivated this research is the degree to which primary sequence alone can stabilize the pre-lasso motif. To conduct an exhaustive sequence exploration, a fixed backbone design was performed with a large subset of class II lasso peptides using Rosetta (51). Comparing different lasso peptides was important due to the diverse structural characteristics exhibited naturally, including variations in sequence length, sizes of the ring, loop, tail, and secondary structural elements (**Figure 1A**). The NMR solution or crystal structures of each peptide were first energy minimized using Rosetta `FastRelax` (56). For each design iteration, all residue positions, excluding those involved in forming the isopeptide bond, were permitted to explore canonical L- and D-amino acid residues except cysteine. Cysteine residues were excluded due to their experimental propensity to form disulfides that could result in undesirable conformational effects. Moreover, if a native sequence featured a bulky aromatic stopper residue (as illustrated in **Figure 2**), the sampling was restricted to Phe, Trp, and Tyr at that specific position. This stopper residue resides beneath the lasso ring, serving as a sterically encumbered plug to prevent unthreading of the C-terminal tail. Following the design phase, each structure underwent scoring using Rosetta’s default score function, REF2015 (75). Rosetta’s REF2015 scoring function is an energy model that evaluates protein structures by accounting for both physical interactions (such as van der Waals forces, hydrogen bonding, and solvation effects) and statistical potentials derived from experimental data. It integrates these contributions to estimate the thermodynamic stability and favorability of a given conformation. The best scoring sequence for each lasso peptide is shown in **Table S1**, sorted by score.

**Figure 2.**
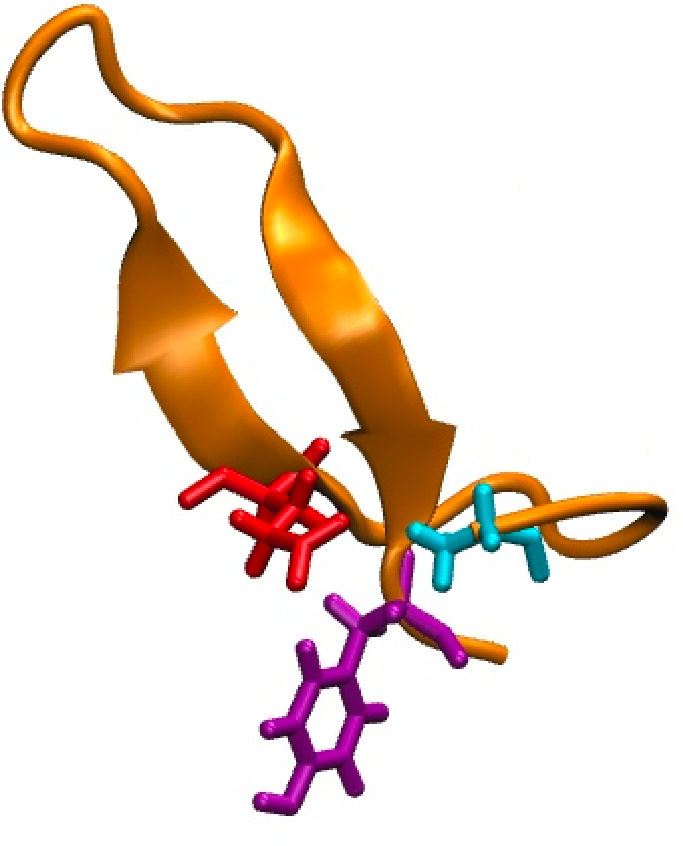
Pre-lasso structure of microcin J25 highlighting the N-terminal and side-chain carboxyl residues involved in isopeptide bond formation and the plug residue (colored cyan, red, and purple, respectively).

Interestingly, fixed-backbone design of these lasso peptides yielded minimal to no sequence variation across the designed sequences for each case. For example, no sequence variation was observed in the designed sequences of MccJ25 (**Figure S1**). To explore this further, additional design runs were performed with four additional MccJ25 structures obtained from a 5000 ns unbiased MD simulation (**Figure S2**). This results in a diverse set of sequences across all five design cases (**Figure S3**), highlighting the significance of lasso backbone conformation in influencing sequence variability during design. However, these designed sequences had worse Rosetta scores. Thus, the best-scoring designs from the NMR spectroscopy solution and crystal structure backbones were selected for further analysis.

### Defining CVs

The four best-scoring designed sequences—which were derived from the backbone structures of rubrivinodin, caulosegnin II, achromonodin-1, and MccJ25 (**Figure 3**)—along with their native counterparts, were then selected for enhanced sampling via WTMetaD to assess their relative pre-lasso stabilities. In our earlier work (38), we identified a protocol for defining CVs that clearly distinguish pre-lasso and pre-tadpole conformations to more efficiently quantify the relative probability of pre-lasso folding. HLDA, a form of supervised machine learning, is first employed to identify a decision boundary that best separates pre-lasso and pre-tadpole conformations. Herein, we additionally included unfolded conformations in the HLDA analysis to maximally separate the pre-lasso regions of phase space from both pre-tadpole and unfolded conformations. Based on previous findings (38), we used the C-alpha distances between residues of the macrocyclic ring and the ring-piercing tail residue as HLDA descriptors (**Figure 4A**). These distances effectively capture the N-terminal region wrapping around the peptide tail, which is crucial for forming the knotted structure. First, each sequence was run in standard MD simulations, starting from a pre-lasso conformation and continuing for 2 μs, to generate structural ensembles for subsequent HLDA analysis. Our previously developed triangulation-based algorithm, LATCHED (37) (see Methods), was then used to identify pre-lasso and pre-tadpole structures. Structures with a distance between the N-terminal nitrogen and the carbonyl carbon of the side chain carboxylate greater than 6 Å were deemed unfolded. Pre-lasso, pre-tadpole, and unfolded conformations obtained from standard MD were then included in the HLDA analysis to determine the optimal linear combination of C-alpha distances for each sequence (**Figures S4-S7**). Notably, both the native and designed caulosegnin II sequences consistently unfolded immediately in multiple MD runs initiated from various pre-lasso conformations, leading to their exclusion from the following free energy assessment due to clear instability. The relatively short lasso loop of caulosegnin II (**Figure 3B**) likely makes it difficult to incorporate sufficient stabilizing interactions to prevent the structure from unfolding.

**Figure 3.**
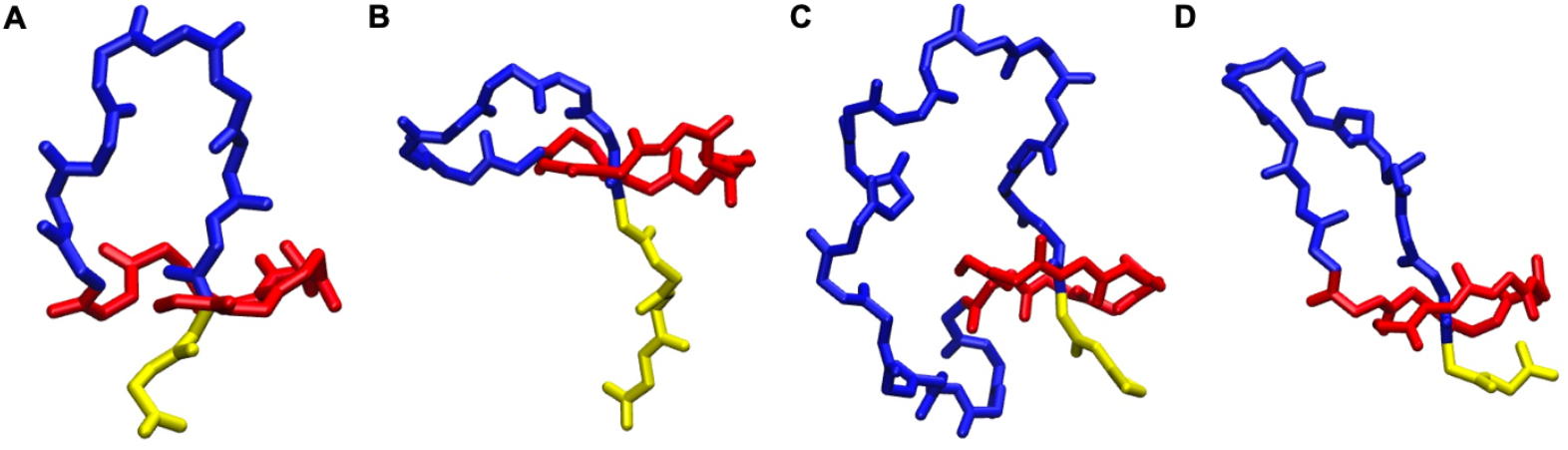
Backbone structures of rubrivinodin (A), caulosegnin II (B), achromonodin-1 (C), and MccJ25 (D). The ring, loop, and tail of each structure are colored red, blue, and yellow, respectively.

**Figure 4.**
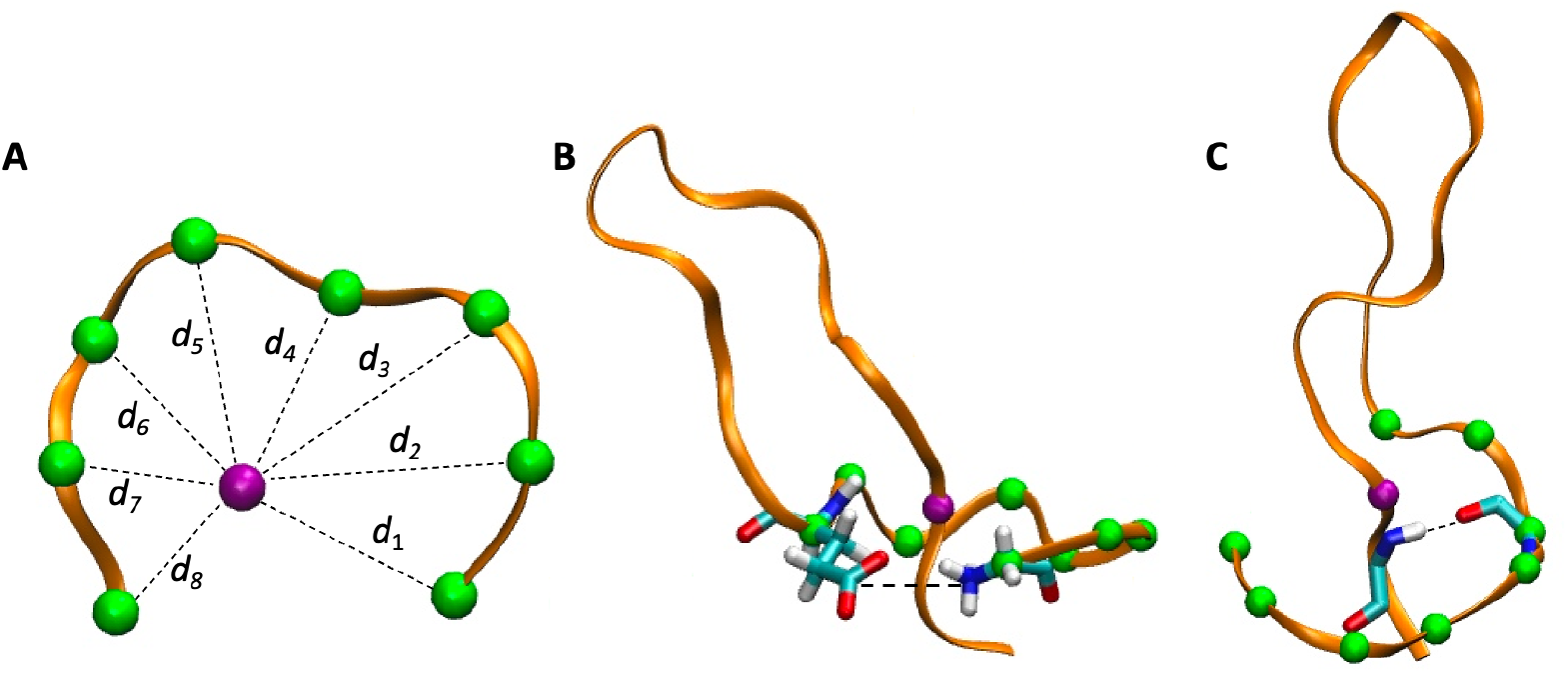
C-alpha distances between residues of the macrocyclic ring (green spheres) and the ring-piercing residue (purple sphere) (A). Coordination between the atoms of the two bond isopeptide-forming residues (B) and atoms of the loop-forming hydrogen bond (C).

In previous work (38), the distance between the two isopeptide bond-forming residues was established as the best CV to use in conjunction with the HLDA CV since the two residues must be proximal during the cyclization of the lasso ring. Although the distance CV was effective in both sampling efficiency and distinguishing pre-lasso and pre-tadpole ensembles, the phase space of the unfolded structures was vast (covering a range between 6-35 Å) and thus dominated the free energy landscape. This disparity makes it difficult to visualize both pre-cyclic and unfolded structures within the same landscape. To improve sampling efficiency and visualization, a coordination number between the two isopeptide bond-forming residues is used herein instead of the distance. The coordination is defined between the N-terminal nitrogen and the carbonyl carbon of the side chain carboxylate (**Figure 4B**) and represented by a switching function at 6 Å in PLUMED to ensure that the calculated CV has continuous derivatives. Using this descriptor instead of the distance between the two atoms focuses the sampling on pre-cyclic structures (with a coordination number > 0) and unfolded structures (coordination of 0) while not focusing sampling time on separating distances beyond the relevant 6 Å pre-cyclic structures. To further increase the separation between the pre-lasso and pre-tadpole + unfolded ensembles, we also used an interaction that occurs upon loop formation (38) (**Figure 4C**). The loop represents the lasso region that is not threaded through the macrocyclic ring (**Figure 1A**) and forms through a turn in the peptide chain of the loop to position the C-terminal tail beneath the encircling ring, thereby forming the hairpin-like structures as seen with MccJ25. This interaction entails a backbone-backbone hydrogen bond between the macrocyclic ring and the first residue of the lasso tail below the ring (residue 6 and 20 for MccJ25, respectively). The hydrogen bond is described here by a coordination number between donor and acceptor atoms with a switching function at 4 Å. A single CV was then defined as the sum of this loop coordination number and the coordination number between the isopeptide bond-forming residues (referred to as Ring + Loop Coordination) to be used in conjunction with the HLDA CV for each sequence.

### Pre-Lasso Stability

Using the HLDA and coordination CVs, the free energy landscapes of native and designed sequences of rubrivinodin, achromonodin-1, and MccJ25 were calculated with WTMetaD (**Figures 5-7 A,B**). Note that the HLDA CV for rubrivinodin flips from being a large number in the native case (**Figure 5A**) to a small number for the designed sequence (**Figure 5B**). We employed our previously developed triangulation-based algorithm, LATCHED (37) (see Methods), to identify pre-lasso and pre-tadpole structures on each landscape, ensuring the separation between the two ensembles (**Figures 5-7 C,D**). The pre-lasso ensemble of each sequence aligns with their respective HLDA CV value (**Figures S1-S4**) and a coordination CV value of ∼1.0 to 2.0 contacts. Regions outside the HLDA CV value for each sequence and shown in red in Figures 5-7 C,D correspond to pre-tadpole structures. All phase space corresponds to unfolded conformations.

**Figure 5.**
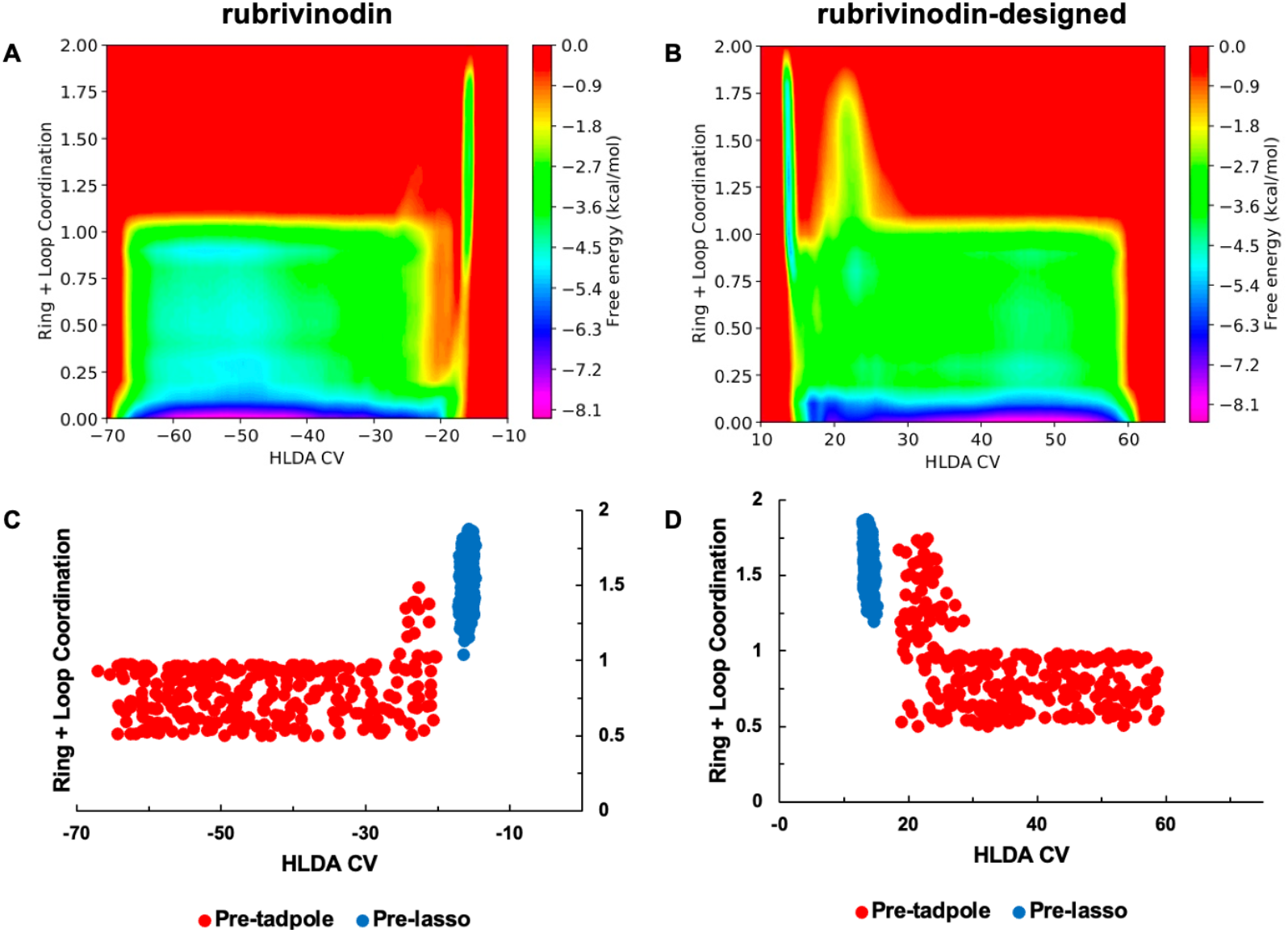
Lasso folding free energy landscape of native (A) and designed (B) sequences of rubrivinodin. The distribution of pre-lasso and pre-tadpole structures for both native (C) and designed (D) sequences is shown.

**Figure 6.**
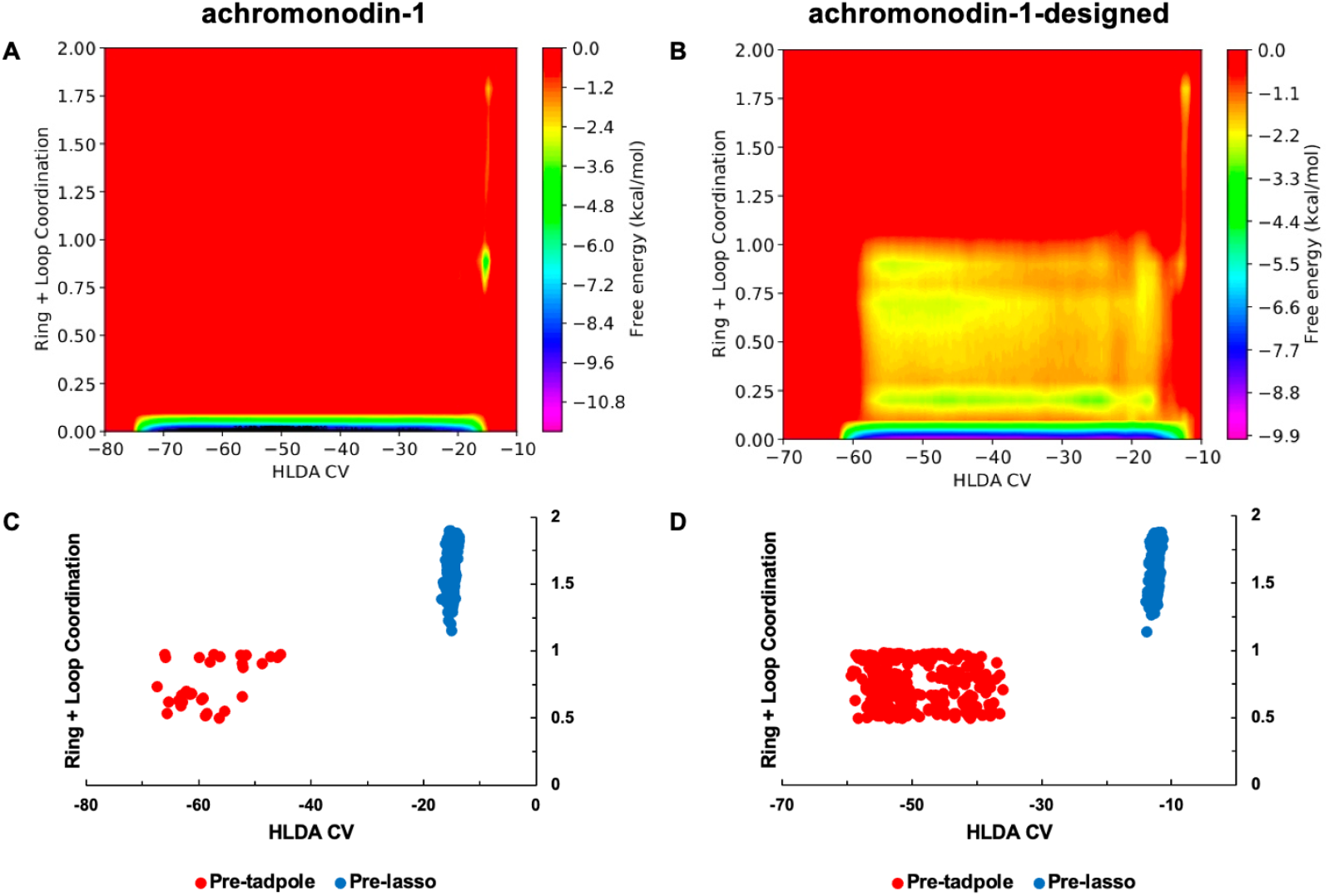
Lasso folding free energy landscape of native (A) and designed (B) sequences of achromonodin-1. The distribution of pre-lasso and pre-tadpole structures for both native (C) and designed (D) sequences is shown.

**Figure 7.**
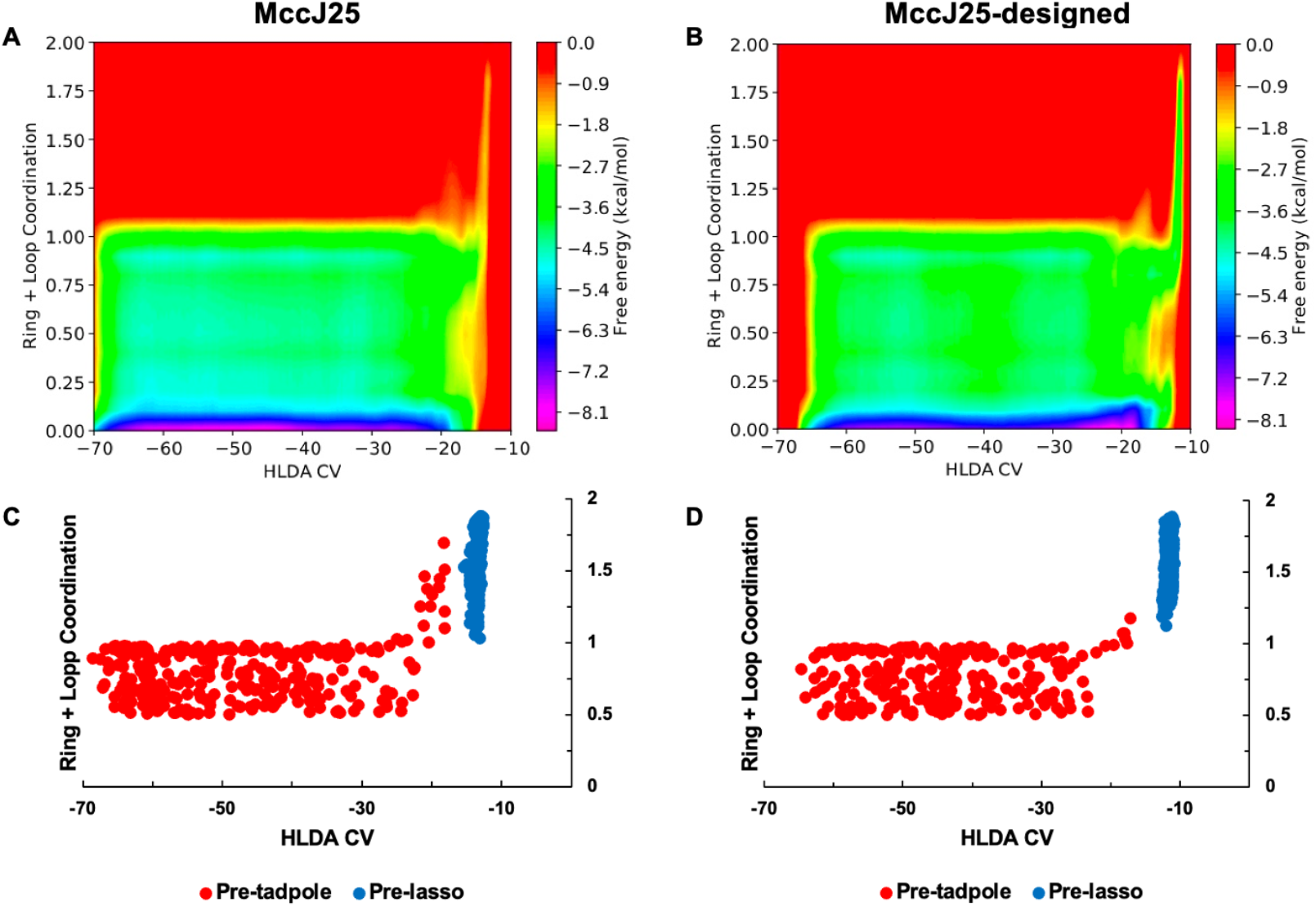
Lasso folding free energy landscape of native (A) and designed (B) sequences of microcin J25. The distribution of pre-lasso and pre-tadpole structures for both native (C) and designed (D) sequences is shown.

By integrating over the pre-lasso and pre-tadpole basins in each landscape, we quantified the relative probability of forming pre-lasso and pre-tadpole structures for each sequence (**Table 1**). Additionally, we normalize these probabilities to pre-cyclic structures and provide a magnitude of pre-lasso gain relative to the native sequence for each case. Assuming the barriers for folding are smaller than those for bond formation such that the equilibrium ratio of pre-lasso and pre-tadpole conformations becomes constant, and the rate of cyclization for these two isomers is not too dissimilar, it is these ratios that are reflective of lasso vs. tadpole cyclization. Sequence design provides a notable increase in the relative pre-lasso stability for rubrivinodin and MccJ25. Both designed sequences showed an increase in the total normalized probability of the pre-lasso state and a slight decrease in the total normalized probability of the pre-tadpole state. For rubrivinodin, the pre-lasso probability normalized to pre-cyclic structures increased from 1.1% for the native sequence to 22.4% for the designed sequence, which was a 20.4-fold improvement in stability. Similarly, for MccJ25 the relative pre-lasso vs. pre-tadpole probability increased from 0.04% for the native sequence to 1.8% for the designed sequence, which was a 45-fold improvement in relative stability. Surprisingly for achromonodin-1, the sequence design slightly increased the probability of the pre-lasso state, but the pre-tadpole state also became more favorable, going from 20.4% to 5.3% pre-lasso to pre-tadpole relative probability, which was a 3.8-fold decrease in stability. These findings indicate that sequence design can provide marked enhancements in pre-lasso stability for some lasso backbone structures, whereas others are more recalcitrant. The backbone arrangements of rubrivinodin and MccJ25 seemingly allow for more favorable interactions to be incorporated via sequence design. This is particularly sensible for MccJ25, which has a beta-hairpin-like secondary structural feature.

**Table 1.**
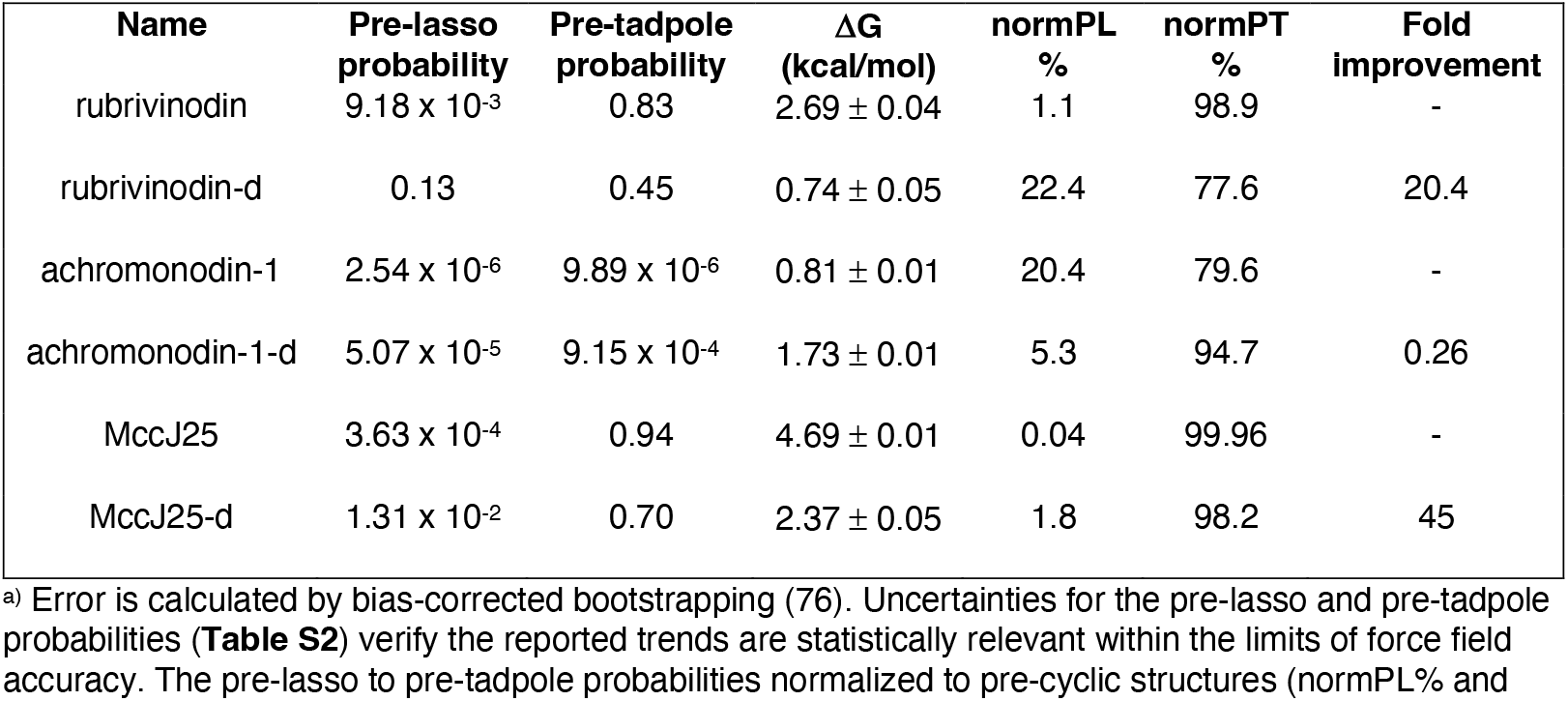
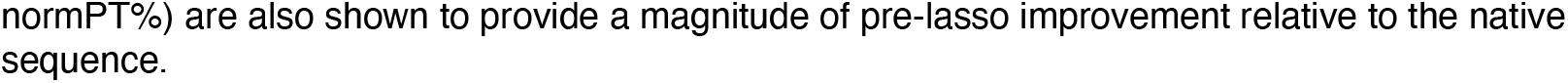
Total and normalized probabilities of forming pre-lasso and pre-tadpole structures, free energy difference between pre-tadpole and pre-lasso (ΔG = G_pre-lasso_ – G_pre-tadpole_)^a^, and fold-improvement of designed sequences (indicated by a -d after the name).

### Chemical modifications

Despite improvements in relative pre-lasso stability for two of the three designed sequences, sequence design alone cannot significantly alter the interaction between the two isopeptide-forming residues (1-8/9 interactions). Additionally, stabilizing this interaction would not only reduce the dominance of unfolded conformations but could also promote pre-lasso stability through ring-loop-tail interactions. This lack of stabilization is partially a consequence of the design strategy since sampling at the positions of the N-terminal and side-chain carboxylate residues was restricted during the design step to preserve the isopeptide bond-forming residues, and therefore no additional interactions could be incorporated to keep the two residues proximal. However, it is also a consequence of steric restriction of the ring wrapping closely around the tail, limiting the two residues from approaching each other. To test the relative impact of modifications to the isopeptide bond-forming residues, moieties used in our earlier work (37) were added to the MccJ25-designed sequence (**Figure 8**). These modifications, capable of chemoselectively forming an isopeptide bond via transthioesterification and S-to-N-acyl transfer events, consist of a thiol containing auxiliary attached to Gly1N (77) and a thioester at position 8 (78). Notably, the thioester used is an Asp-based variant, replacing Glu at position 8. With one less methylene group than Glu, the Asp-based variant offers a closer comparison to the pre-ring size of native pre-MccJ25. For the native MccJ25 (37), these modifications increased the percentage of pre-lasso structures through pi-pi stacking interactions that orient the hydrophobic ring faces and solvated polar groups such that spontaneous unfolding is disfavored (**Figure 8**). Consistent with the native sequence, these additions increase the relative pre-lasso stability of the designed MccJ25 (**Figure 9** and **Table 2**). Notably, when introducing the modifications, a new pre-tadpole conformation that closely resembles a pre-lasso is sampled—making separation between the two ensembles more difficult. However, sufficient separation is still achieved to distinguish pre-lasso from pre-tadpole (**Figure S8**). A significant increase in the probability of both pre-lasso and pre-tadpole structures is observed. This leads to an increase in the normalized pre-lasso probability from 1.8% to 14.1% for the modified MccJ25 designed sequence. This corresponds to a 7.8-fold increase in the stability of the design, and a 352.5-fold increase in stability relative to the native sequence. When the interaction between the bond-forming residues is weak, there is little to offset the entropic freedom of ring fluctuations and increases the likelihood of unfolding. Therefore, we anticipate that a combination of sequence design and non-canonical residue modifications to increase the ring residue interactions will play an essential role in further improving pre-lasso stabilization.

**Figure 8.**
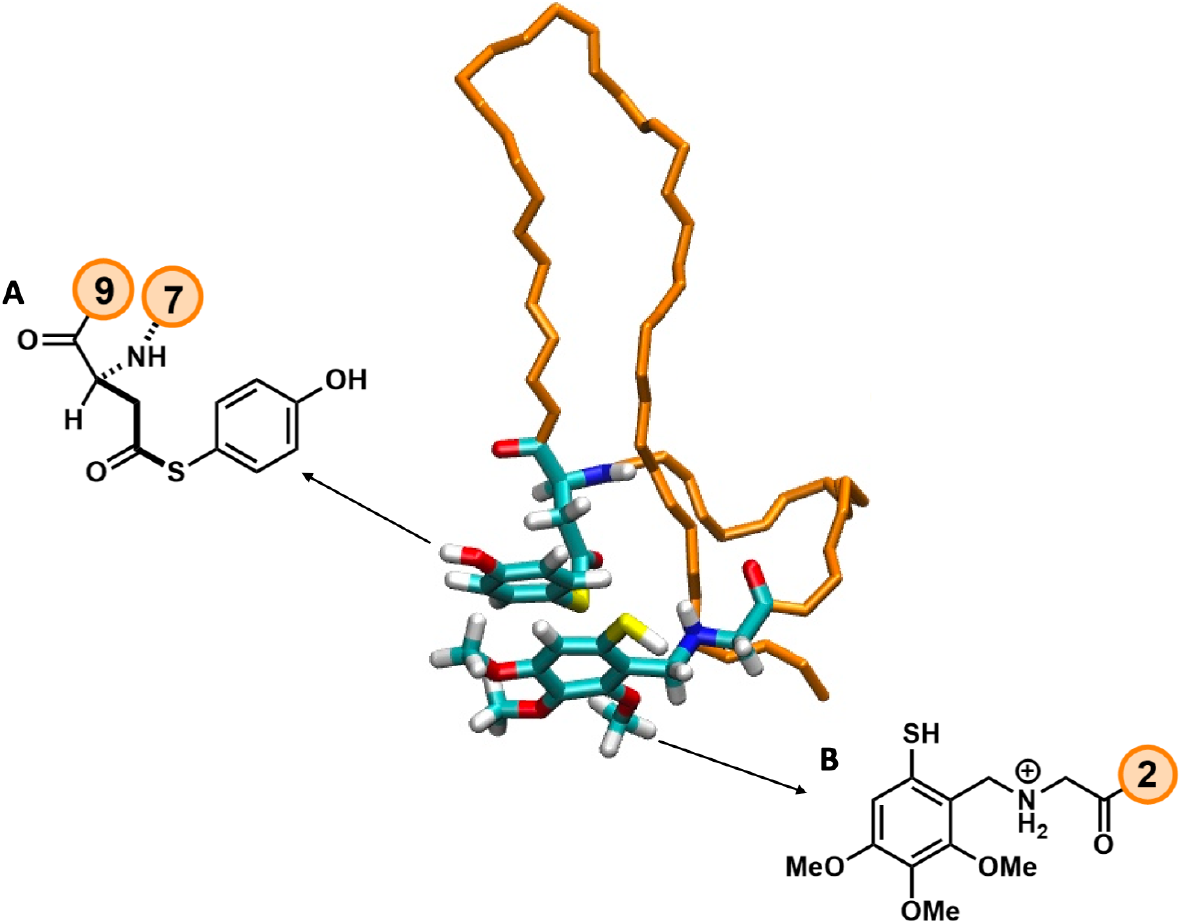
Structures of the residue modifications added to the N-terminal residue (B) and the side-chain carboxyl (A) residues of MccJ25-designed.

**Figure 9.**
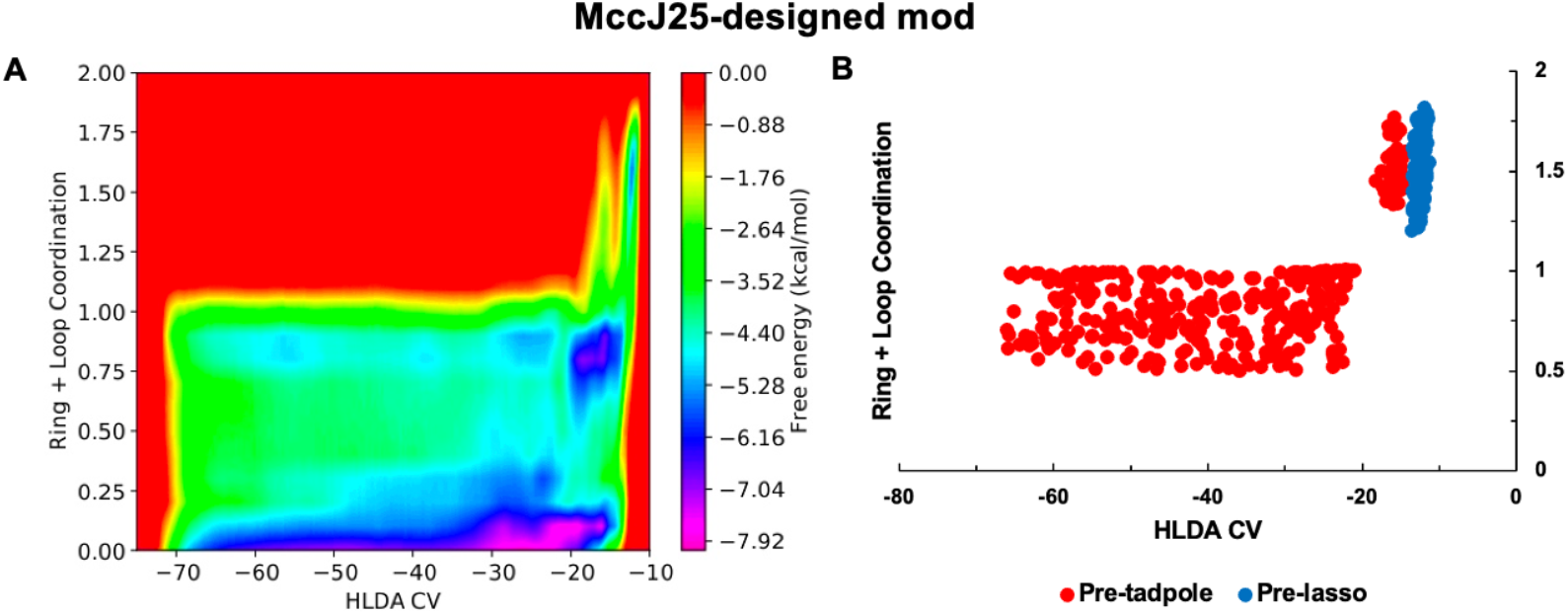
Lasso folding free energy landscape of the microcin J25-designed sequence with chemical modifications (A) and the distribution of pre-lasso and pre-tadpole structures (B).

**Table 2.** Total and normalized probabilities of forming pre-lasso and pre-tadpole structures, free energy difference between pre-tadpole and pre-lasso (ΔG = G_pre-lasso_ – G_pre-tadpole_)^a^, and fold-improvement of designed (-d), modified (mod) and disulfide-linked sequences.

| Name | Pre-lasso probability | Pre-tadpole probability | $\Delta G$ (kcal/mol) | normPL% | normPT % | Fold improvement |
| --- | --- | --- | --- | --- | --- | --- |
| MccJ25 | $3.63 \times 10^{-4}$ | 0.94 | $4.69 \pm 0.01$ | 0.04 | 99.96 | - |
| MccJ25-d | $1.31 \times 10^{-2}$ | 0.70 | $2.37 \pm 0.05$ | 1.8 | 98.2 | 45 |
| MccJ25-d mod | 1.40 | 8.53 | $1.08 \pm 0.13$ | 14.1 | 85.9 | 352.5 |
| MccJ25-d disulfide | $9.56 \times 10^{-4}$ | 2.01 | $4.56 \pm 0.03$ | $4.75 \times 10^{-2}$ | 99.95 | 1.2 |
| MccJ25-d mod + disulfide | 3.26 | 12.57 | $0.80 \pm 0.12$ | 20.6 | 79.4 | 515 |

<sup>a)</sup> Error is calculated by bias-corrected bootstrapping (76). Uncertainties for the pre-lasso and pre-tadpole probabilities are presented in **Table S2**. The pre-lasso to pre-tadpole probabilities normalized to pre-cyclic structures (normPL% and normPT%) are also shown to provide a magnitude of pre-lasso accessibility relative to the native, unmodified sequence.

### Cross-Linking to Decrease Entropic Freedom

Examining the free energy landscapes obtained thus far, a common feature is the vastness of the unfolded and pre-tadpole conformation space relative to the pre-lasso, making it difficult to access the pre-lasso from the unfolded and pre-tadpole ensembles. This is supported by our analysis of the relative entropy of native MccJ25 in our previous study (37). One way to potentially reduce the unfolded and pre-tadpole space is by incorporating a cross-link. Cross-links have been used to constrain peptides by stabilizing the structures of alpha-helices and beta-hairpins (79,80). To incorporate a cross-link into a lasso peptide, we used Rosetta `Disulfidize` (81) to identify and build potential L and D-cysteine disulfides in the native lasso structures of rubrivinodin, achromonodin-1, and MccJ25. The analysis was done for two sets of residues in each lasso sequence: 1) residues of the macrocyclic ring, and 2) residues of the loop and tail regions. These two sets of residues were selected because a cross-link connecting them would create the most conformationally restricted lasso structure, akin to Class I and III lassos. This resulted in one identified disulfide between residues Cys7 and Cys16 of MccJ25-designed, connecting the ring to the loop (**Figure 10**). A second `Disulfidize` analysis using residues of the loop of rubrivinodin, achromonodin-1, and MccJ25 was performed. However, no additional disulfide placements were found. The identification of only one disulfide bond in the analysis is likely due to the compact, knot-like structure of the lasso motif, which limits the available space between residues for potential disulfide cross-links. Fixed backbone design was then performed with the MccJ25-disulfide structure to optimize the sequence around the disulfide using the same design protocol described above. Interestingly, this resulted in the same MccJ25-designed sequence excluding the required Cys7 and Cys16 residues. Evaluating the cross-linked sequence with WTMetaD, we found that the addition of the disulfide removed a significant amount of unfolded and pre-tadpole phase space in the free energy landscape (**Figure 11**). Relative to the landscape of the MccJ25-designed sequence (**Figure 7B**), the HLDA space corresponding to unfolded and pre-tadpole structures was reduced by ∼15 units.

**Figure 10.**
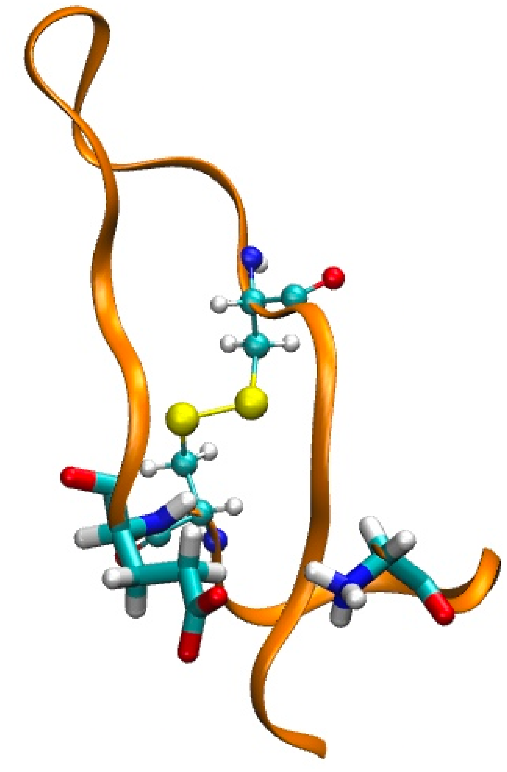
Structure of MccJ25-designed with a disulfide connecting residues 7 and 16. The isopeptide bond-forming residues are shown as licorice sticks while residues 7 and 16 are depicted as ball-stick representation.

**Figure 11.**
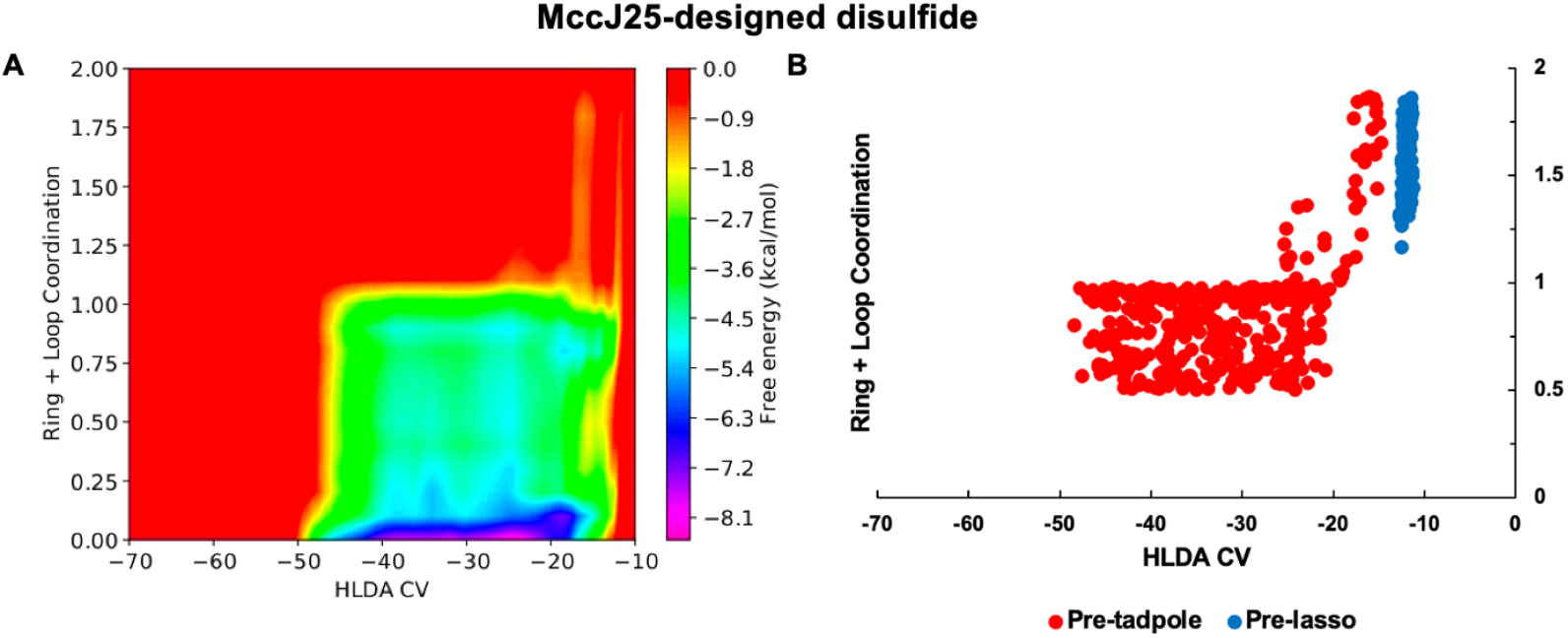
Lasso folding free energy landscape of the microcin J25-designed sequence with a disulfide cross-link (A) and the distribution of pre-lasso and pre-tadpole structures (B).

However, despite this reduction in phase space, the addition of the disulfide significantly decreased the probability of the pre-lasso state and increased the probability of the pre-tadpole state—resulting in only a minimal increase in the normalized pre-lasso probability of 0.04% to 0.05% compared to the native sequence, but a 36-fold decrease in stability compared to the design without the disulfide (**Table 2**). Although the connection between residues 7 and 16 prevents the loop from completely unraveling, the added constraint to the ring that the disulfide imparts likely makes it more difficult for the N-terminal end to wrap around the C-terminal tail and form the pre-lasso ring, indirectly favoring more pre-tadpole conformations.

Finally, we added the modifications in **Figure 8** to the cross-linked sequence to finish our evaluation of strategies to increase relative pre-lasso stability. We found that the modifications paired with the disulfide cross-link significantly increases the probability of both pre-lasso and pre-tadpole states (**Figure 12** and **Table 2**). The modified designed MccJ25 sequence goes from a normalized pre-lasso probability of 14.1% to 20.6% with the disulfide (1.5-fold increase) and gives a 515-fold increase in stability relative to the native sequence. The enlargement of the pre-lasso ring due to the introduction of the modifications likely counteracts some of the strain imparted by the cross-link when the ring wraps around the tail, which also combines with the stabilizing effects that result from the interaction of the modifications to achieve the highest degree of stabilization relative to the native sequence. Taken together, the results in **Table 2** indicate that the addition of modifications can be highly effective in stabilizing the pre-lasso state, while incorporating a cross-link can be counterproductive if the imparted constraint is not counteracted.

**Figure 12.**
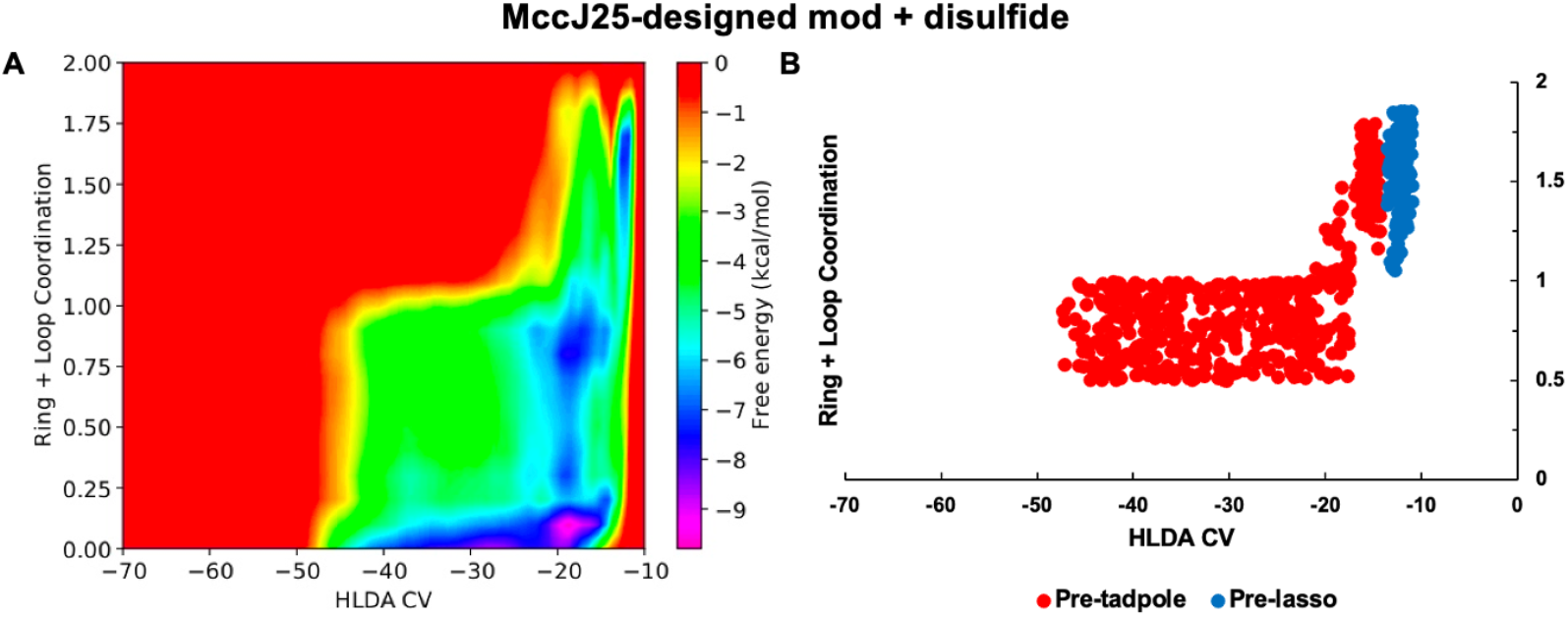
Lasso folding free energy landscape of the microcin J25-designed sequence with chemical modifications and a disulfide cross-link (A). The distribution of pre-lasso and pre-tadpole structures is shown (B).\

## CONCLUSIONS

This work explores effective design strategies for stabilizing de novo lasso peptide folding. Focusing on the relative stability of the pre-lasso intermediate that leads to the interlocked cyclic motif, we test the impacts of optimizing the primary sequence, including non-canonical amino acid residues to stabilize pre-ring formation, and cross-linking the peptide to decrease the entropic dominance of the unfolded and unthreaded ensembles. Optimization of primary sequence through multiple rounds of Rosetta fixed-backbone design is shown to improve the pre-lasso stability for the rubrivinodin and MccJ25 lasso peptides relative to their native counterparts. However, caulosegnin II and achromonodin-1 proved challenging to stabilize, suggesting that the native backbone configuration could play an integral role in the potential peptide stabilization. Moreover, the marginally stable designed sequence of MccJ25 can be considerably increased by incorporating modifications to the isopeptide bond-forming residues. In contrast, including a disulfide cross-link to MccJ25 adversely impacted stability. However, cross-linking combined with bond-forming modifications resulted in the most significant improvement relative to the native sequence. Collectively, sequence optimization, chemical modification, and cross-linking stabilize the pre-lasso motif over 515-fold, such that it represents >20% of the pre-cyclic structures required to form the isopeptide bond. This suggests that this type of combined strategy could enable synthetic access to lasso peptides through sufficient introduction of stabilizing components such as the final MccJ25 case. Future work will continue to investigate alternative potential chemical modifications and cross-links.

Regarding design strategies, we find that fixed-backbone design on NMR spectroscopy solution or crystal structures provides a diverse set of sequences with varied Rosetta scores across the range of lasso peptides tested. Since two of the top four scoring sequences showed improved relative pre-lasso stability when the free energies were evaluated with WTMetaD, this is indeed a useful design method. However, the other two top-scoring sequences did not show stability improvement in free energy analysis. Moreover, these relative trends could not be deciphered from short, unbiased MD runs following design. Only by conducting a free energy evaluation was stabilization verified. This also suggests that free energy analysis of lower-scoring designed sequences could provide additional promising lasso sequences. A free energy analysis is also likely required to accurately assess different design strategies, such as fixed-backbone design vs. flexible backbone design and designing with lasso structures vs. pre-lassos. A limitation of fixed-backbone design used herein was its inability to generate sequence variation for a given lasso peptide. Thus, future efforts could incorporate conformational backbone diversity to increase sequence variability. Additionally, lasso peptides lacking NMR spectroscopy solution or crystal structures could be considered for design by employing the recently developed LassoPred structure prediction tools specifically designed for lasso peptides, and hosting a database of 13,866 structures obtained from 4,749 predicted natural lasso sequences (40).

Moreover, the structure prediction tool has been extended to predict unnatural left-handed lasso peptide structures. Design efforts could also explore cases where the lasso conformation is highly adaptable. For example, lasso peptides such as benenodin-1 undergo conformational switching between two distinct lasso conformers upon heating (41), a process involving extensive restructuring of intramolecular non-covalent interactions (82). Designing sequences that optimize both conformers could not only provide new variants but also enhance the conformational flexibility, and any potentially related function of the lasso structure. Finally, considering the extensive simulation time required (33.6 μs) to evaluate the relative pre-lasso stability of just three designed sequences and their native counterparts, it is essential to develop a streamlined workflow. Ideally, such a workflow should automate the following steps for each sequence in parallel: 1) running standard MD, 2) performing HLDA to obtain CVs, 3) running WTMetaD and verifying convergence, and 4) calculating normalized probabilities. While significantly slower than Rosetta design alone, this automation will increase the overall efficiency, while retaining the fidelity of the relative accessibility of the pre-cyclic conformers.

Collectively, this work provides strategies and suggestions for the continued development of accessing lasso peptides via chemical synthesis and de novo folding. The physical constraints of de novo folding are clearly different from enzyme-stabilized folding. The latter has recently emerged as a powerful tool for expanding lasso sequence diversification through the development of novel biosynthetic strategies, computationally guided lasso cyclase engineering, and models for substrate compatibility prediction. Similarly, we hope that continued development of a streamlined workflow to identify the stabilization criteria underlying *de novo* folding design principles combined with tunable synthetic approaches for introducing backbone modifications, cross-linking, and a compatible cyclization strategy could enable expanded sequence diversity potentially accessible via chemical synthesis. This approach would provide a complimentary platform to biosynthetic methods for accessing lasso peptides with sequences that include non-canonical residues (22,26,83), as well as thus far inaccessible peptidomimetic components and left-handed structures, thereby contributing additional tools for diversified access to these remarkable scaffolds to expand their therapeutic and biotechnological potential.

## SUPPORTING MATERIAL

Supporting Material can be found online at https://doi.org/XX.YYYY/j.bpj.2024.0Z.ZZZ

## AUTHOR CONTRIBUTIONS

All authors designed the research. JDMN and GCAH performed the simulations and analyses. All authors interpreted the results and wrote the manuscript.

## ACKNOWLEDGEMENTS

We thank Dr. Andrew Watkins, Director of Machine Learning for Drug Discovery, Genentech Research and Early Development, for helpful guidance on Rosetta design strategies. JDMN, GCAH, and JMJS acknowledge support from NIH NIGMS (R35GM143117) and computational resources provided by Bridges-2 at the Pittsburgh Supercomputing Center through the ACCESS program (allocation MCB200018) supported by NSF (grants #2138259, #2138286, #2138307, #2137603, and #2138296), as well as the Center for High-Performance Computing (CHPC) at the University of Utah. MCM and AGR acknowledge support from NIH NIGMS (R35GM155457).

## REFERENCES

1. Ongpipattanakul, C., E. K. Desormeaux, A. DiCaprio, W. A. van der Donk, D. A. Mitchell, and S. K. Nair. 2022. Mechanism of Action of Ribosomally Synthesized and Post-Translationally Modified Peptides. Chem Rev. 122(18):14722–14814, doi: 10.1021/acs.chemrev.2c00210.

2. Potterat, O., K. Wagner, G. Gemmecker, J. Mack, C. Puder, R. Vettermann, and R. Streicher. 2004. BI-32169, a Bicyclic 19-Peptide with Strong Glucagon Receptor Antagonist Activity from Streptomyces sp. Journal of Natural Products. 67(9):1528–1531, doi: 10.1021/np040093o.

3. Salomón, R. A., and R. N. Farías. 1992. Microcin 25, a novel antimicrobial peptide produced by Escherichia coli. J Bacteriol. 174(22):7428–7435, doi: 10.1128/jb.174.22.7428-7435.1992.

4. Kodani, S., and K. Unno. 2020. How to harness biosynthetic gene clusters of lasso peptides. J Ind Microbiol Biotechnol. 47(9-10):703–714, doi: 10.1007/s10295-020-02292-6.

5. Martin-Gomez, H., and J. Tulla-Puche. 2018. Lasso peptides: chemical approaches and structural elucidation. Org Biomol Chem. 16(28):5065–5080, doi: 10.1039/c8ob01304g.

6. Ferguson, A. L., S. Zhang, I. Dikiy, A. Z. Panagiotopoulos, P. G. Debenedetti, and A. James Link. 2010. An experimental and computational investigation of spontaneous lasso formation in microcin J25. Biophys J. 99(9):3056–3065, doi: 10.1016/j.bpj.2010.08.073.

7. Rosengren, K. J., R. J. Clark, N. L. Daly, U. Göransson, A. Jones, and D. J. Craik. 2003. Microcin J25 Has a Threaded Sidechain-to-Backbone Ring Structure and Not a Head-to-Tail Cyclized Backbone. Journal of the American Chemical Society. 125(41):12464–12474, doi: 10.1021/ja0367703.

8. Lear, S., T. Munshi, A. S. Hudson, C. Hatton, J. Clardy, J. A. Mosely, T. J. Bull, C. S. Sit, and S. L. Cobb. 2016. Total chemical synthesis of lassomycin and lassomycin-amide. Org Biomol Chem. 14(19):4534–4541, doi: 10.1039/c6ob00631k.

9. Waliczek, M., M. Wierzbicka, M. Arkuszewski, M. Kijewska, L. Jaremko, P. Rajagopal, K. Szczepski, A. Sroczynska, M. Jaremko, and P. Stefanowicz. 2020. Attempting to synthesize lasso peptides using high pressure. PLoS One. 15(6):e0234901, doi: 10.1371/journal.pone.0234901.

10. Hegemann, J. D. 2020. Factors Governing the Thermal Stability of Lasso Peptides. Chembiochem. 21(1-2):7–18, doi: 10.1002/cbic.201900364.

11. Duquesne, S., D. Destoumieux-Garzon, S. Zirah, C. Goulard, J. Peduzzi, and S. Rebuffat. 2007. Two enzymes catalyze the maturation of a lasso peptide in Escherichia coli. Chem Biol. 14(7):793–803, doi: 10.1016/j.chembiol.2007.06.004.

12. Hegemann, J. D., M. Zimmermann, X. Xie, and M. A. Marahiel. 2015. Lasso Peptides: An Intriguing Class of Bacterial Natural Products. Accounts of Chemical Research. 48(7):1909–1919, doi: 10.1021/acs.accounts.5b00156.

13. Barrett, S. E., S. Yin, P. Jordan, J. K. Brunson, J. Gordon-Nunez, G. Costa Machado da Cruz, C. Rosario, B. K. Okada, K. Anderson, T. A. Pires, R. Wang, D. Shukla, M. J. Burk, and D. A. Mitchell. 2024. Substrate interactions guide cyclase engineering and lasso peptide diversification. Nat Chem Biol. doi: 10.1038/s41589-024-01727-w.

14. Hegemann, J. D., M. Zimmermann, X. Xie, and M. A. Marahiel. 2013. Caulosegnins I-III: a highly diverse group of lasso peptides derived from a single biosynthetic gene cluster. J Am Chem Soc. 135(1):210–222, doi: 10.1021/ja308173b.

15. Ducasse, R., K. P. Yan, C. Goulard, A. Blond, Y. Li, E. Lescop, E. Guittet, S. Rebuffat, and S. Zirah. 2012. Sequence determinants governing the topology and biological activity of a lasso peptide, microcin J25. Chembiochem. 13(3):371–380, doi: 10.1002/cbic.201100702.

16. Si, Y., A. M. Kretsch, L. M. Daigh, M. J. Burk, and D. A. Mitchell. 2021. Cell-Free Biosynthesis to Evaluate Lasso Peptide Formation and Enzyme-Substrate Tolerance. J Am Chem Soc. 143(15):5917–5927, doi: 10.1021/jacs.1c01452.

17. DiCaprio, A. J., A. Firouzbakht, G. A. Hudson, and D. A. Mitchell. 2019. Enzymatic Reconstitution and Biosynthetic Investigation of the Lasso Peptide Fusilassin. J Am Chem Soc. 141(1):290–297, doi: 10.1021/jacs.8b09928.

18. Hills, E., T. J. Woodward, S. Fields, and B. M. Brandsen. 2022. Comprehensive Mutational Analysis of the Lasso Peptide Klebsidin. ACS Chem Biol. 17(4):998–1010, doi: 10.1021/acschembio.2c00148.

19. Liu, T., X. Ma, J. Yu, W. Yang, G. Wang, Z. Wang, Y. Ge, J. Song, H. Han, W. Zhang, D. Yang, X. Liu, and M. Ma. 2021. Rational generation of lasso peptides based on biosynthetic gene mutations and site-selective chemical modifications. Chem Sci. 12(37):12353–12364, doi: 10.1039/d1sc02695j.

20. Fernandez, H. N., A. M. Kretsch, S. Kunakom, A. E. Kadjo, D. A. Mitchell, and A. S. Eustaquio. 2024. High-Yield Lasso Peptide Production in a Burkholderia Bacterial Host by Plasmid Copy Number Engineering. ACS Synth Biol. 13(1):337–350, doi: 10.1021/acssynbio.3c00597.

21. Thokkadam, A., T. Do, X. Ran, M. P. Brynildsen, Z. J. Yang, and A. J. Link. 2023. High-Throughput Screen Reveals the Structure-Activity Relationship of the Antimicrobial Lasso Peptide Ubonodin. ACS Cent Sci. 9(3):540–550, doi: 10.1021/acscentsci.2c01487.

22. Schiefelbein, K., J. Lang, M. Schuster, C. E. Grigglestone, R. Striga, L. Bigler, M. C. Schuman, O. Zerbe, Y. Li, and N. Hartrampf. 2024. Merging Flow Synthesis and Enzymatic Maturation to Expand the Chemical Space of Lasso Peptides. J Am Chem Soc. 146(25):17261–17269, doi: 10.1021/jacs.4c03898.

23. Cheung, W. L., M. Y. Chen, M. O. Maksimov, and A. J. Link. 2016. Lasso Peptide Biosynthetic Protein LarB1 Binds Both Leader and Core Peptide Regions of the Precursor Protein LarA. ACS Cent Sci. 2(10):702–709, doi: 10.1021/acscentsci.6b00184.

24. Pan, S. J., and A. J. Link. 2011. Sequence diversity in the lasso peptide framework: discovery of functional microcin J25 variants with multiple amino acid substitutions. J Am Chem Soc. 133(13):5016–5023, doi: 10.1021/ja1109634.

25. Knappe, T. A., F. Manzenrieder, C. Mas-Moruno, U. Linne, F. Sasse, H. Kessler, X. Xie, and M. A. Marahiel. 2011. Introducing Lasso Peptides as Molecular Scaffolds for Drug Design: Engineering of an Integrin Antagonist. Angewandte Chemie International Edition. 50(37):8714–8717, doi: 10.1002/anie.201102190.

26. Piscotta, F. J., J. M. Tharp, W. R. Liu, and A. J. Link. 2015. Expanding the chemical diversity of lasso peptide MccJ25 with genetically encoded noncanonical amino acids. Chemical Communications. 51(2):409–412, doi: 10.1039/C4CC07778D.

27. Schroder, H. V., Y. Zhang, and A. J. Link. 2021. Dynamic covalent self-assembly of mechanically interlocked molecules solely made from peptides. Nat Chem. 13(9):850–857, doi: 10.1038/s41557-021-00770-7.

28. Baquero, F., K. Beis, D. J. Craik, Y. Li, A. J. Link, S. Rebuffat, R. Salomon, K. Severinov, S. Zirah, and J. D. Hegemann. 2024. The pearl jubilee of microcin J25: thirty years of research on an exceptional lasso peptide. Nat Prod Rep. 41(3):469–511, doi: 10.1039/d3np00046j.

29. Tietz, J. I., C. J. Schwalen, P. S. Patel, T. Maxson, P. M. Blair, H. C. Tai, U. I. Zakai, and D. A. Mitchell. 2017. A new genome-mining tool redefines the lasso peptide biosynthetic landscape. Nat Chem Biol. 13(5):470–478, doi: 10.1038/nchembio.2319.

30. Mi, X., S. E. Barrett, D. A. Mitchell, D. Shukla, and 2024. LassoESM: A tailored language model for enhanced lasso peptide property prediction. bioRxiv. doi: 10.1101/2024.10.25.620295.

31. Digal, L., S. C. Samson, M. A. Stevens, A. Ghorai, H. Kim, M. C. Mifflin, K. R. Carney, D. L. Williamson, S. Um, G. Nagy, D. C. Oh, M. C. Mendoza, and A. G. Roberts. 2024. Nonthreaded Isomers of Sungsanpin and Ulleungdin Lasso Peptides Inhibit H1299 Cancer Cell Migration. ACS Chem Biol. 19(1):81–88, doi: 10.1021/acschembio.3c00525.

32. Wilson, K.-A., M. Kalkum, J. Ottesen, J. Yuzenkova, B. T. Chait, R. Landick, T. Muir, K. Severinov, and S. A. Darst. 2003. Structure of Microcin J25, a Peptide Inhibitor of Bacterial RNA Polymerase, is a Lassoed Tail. Journal of the American Chemical Society. 125(41):12475–12483, doi: 10.1021/ja036756q.

33. Soudy, R., L. Wang, and K. Kaur. 2012. Synthetic peptides derived from the sequence of a lasso peptide microcin J25 show antibacterial activity. Bioorg Med Chem. 20(5):1794–1800, doi: 10.1016/j.bmc.2011.12.061.

34. Hammami, R., F. Bedard, A. Gomaa, M. Subirade, E. Biron, and I. Fliss. 2015. Lasso-inspired peptides with distinct antibacterial mechanisms. Amino Acids. 47(2):417–428, doi: 10.1007/s00726-014-1877-x.

35. Katahira, R., K. Shibata, M. Yamasaki, Y. Matsuda, and M. Yoshida. 1995. RES-701-1, comparative study of the synthetic and the microbial-origin compounds. Bioorg. Med. Chem. Lett. 5(15):1595–1600, doi: 10.1016/0960894X9500264T.

36. Ferguson, A. L., S. Zhang, I. Dikiy, A. Z. Panagiotopoulos, P. G. Debenedetti, and A. J. Link. 2010. An experimental and computational investigation of spontaneous lasso formation in microcin J25. Biophysical Journal. 99(9):3056–3065, doi: 10.1016/j.bpj.2010.08.073.

37. da Hora, G. C. A., M. Oh, M. C. Mifflin, L. Digal, A. G. Roberts, and J. M. J. Swanson. 2024. Lasso Peptides: Exploring the Folding Landscape of Nature’s Smallest Interlocked Motifs. J Am Chem Soc. 146(7):4444–4454, doi: 10.1021/jacs.3c10126.

38. da Hora, G. C. A., M. Oh, J. D. M. Nguyen, and J. M. J. Swanson. 2024. One Descriptor to Fold Them All: Harnessing Intuition and Machine Learning to Identify Transferable Lasso Peptide Reaction Coordinates. J Phys Chem B. 128(17):4063–4075, doi: 10.1021/acs.jpcb.3c08492.

39. Juarez, R. J., Y. Jiang, M. Tremblay, Q. Shao, A. J. Link, and Z. J. Yang. 2023. LassoHTP: A High-Throughput Computational Tool for Lasso Peptide Structure Construction and Modeling. J Chem Inf Model. 63(2):522–530, doi: 10.1021/acs.jcim.2c00945.

40. Ouyang, X., X. Ran, H. Xu, Y. Zhao, A. J. Link, and Z. Yang. 2024. Predicting 3D Structures of Lasso Peptides. ChemRxiv. doi: 10.26434/chemrxiv-2024-q3rn0-v2.

41. Yang, Z., N. Hajlasz, and H. J. Kulik. 2022. Computational Modeling of Conformer Stability in Benenodin-1, a Thermally Actuated Lasso Peptide Switch. The Journal of Physical Chemistry B. 126(18):3398–3406, doi: 10.1021/acs.jpcb.2c00762.

42. Lai, P.-K., and Y. N. Kaznessis. 2017. Free Energy Calculations of Microcin J25 Variants Binding to the FhuA Receptor. Journal of Chemical Theory and Computation. 13(7):3413–3423, doi: 10.1021/acs.jctc.7b00417.

43. Allen, C. D., M. Y. Chen, A. Y. Trick, D. T. Le, A. L. Ferguson, and A. J. Link. 2016. Thermal Unthreading of the Lasso Peptides Astexin-2 and Astexin-3. ACS Chem Biol. 11(11):3043–3051, doi: 10.1021/acschembio.6b00588.

44. Tan, H. N., W. Q. Liu, J. Ho, Y. J. Chen, F. J. Shieh, H. T. Liao, S. P. Wang, J. D. Hegemann, C. Y. Chang, and J. Chu. 2024. Structure Prediction and Protein Engineering Yield New Insights into Microcin J25 Precursor Recognition. ACS Chem Biol. 19(9):1982–1990, doi: 10.1021/acschembio.4c00251.

45. Yao, G., S. Kosol, M. T. Wenz, E. Irran, B. G. Keller, O. Trapp, and R. D. Sussmuth. 2022. The occurrence of ansamers in the synthesis of cyclic peptides. Nat Commun. 13(1):6488, doi: 10.1038/s41467-022-34125-8.

46. Schuler, L. D., and W. F. Van Gunsteren. 2000. On the Choice of Dihedral Angle Potential Energy Functions for n-Alkanes. Molecular Simulation. 25(5):301–319, doi: 10.1080/08927020008024504.

47. Schuler, L. D., X. Daura, and W. F. van Gunsteren. 2001. An improved GROMOS96 force field for aliphatic hydrocarbons in the condensed phase. Journal of Computational Chemistry. 22(11):1205–1218, doi: 10.1002/jcc.1078.

48. Oostenbrink, C., T. A. Soares, N. F. A. van der Vegt, and W. F. van Gunsteren. 2005. Validation of the 53A6 GROMOS force field. European Biophysics Journal : EBJ. 34(4):273–284, doi: 10.1007/s00249-004-0448-6.

49. Schmid, N., A. P. Eichenberger, A. Choutko, S. Riniker, M. Winger, A. E. Mark, and W. F. van Gunsteren. 2011. Definition and testing of the GROMOS force-field versions 54A7 and 54B7. Eur Biophys J. 40(7):843–856, doi: 10.1007/s00249-011-0700-9.

50. Soares, T. A., X. Daura, C. Oostenbrink, L. J. Smith, and W. F. van Gunsteren. 2004. Validation of the GROMOS force-field parameter set 45A3 against nuclear magnetic resonance data of hen egg lysozyme. Journal of Biomolecular NMR. 30(4):407–422, doi: 10.1007/s10858-004-5430-1.

51. Leman, J. K., B. D. Weitzner, S. M. Lewis, J. Adolf-Bryfogle, N. Alam, R. F. Alford, M. Aprahamian, D. Baker, K. A. Barlow, P. Barth, B. Basanta, B. J. Bender, K. Blacklock, J. Bonet, S. E. Boyken, P. Bradley, C. Bystroff, P. Conway, S. Cooper, B. E. Correia, B. Coventry, R. Das, R. M. De Jong, F. DiMaio, L. Dsilva, R. Dunbrack, A. S. Ford, B. Frenz, D. Y. Fu, C. Geniesse, L. Goldschmidt, R. Gowthaman, J. J. Gray, D. Gront, S. Guffy, S. Horowitz, P. S. Huang, T. Huber, T. M. Jacobs, J. R. Jeliazkov, D. K. Johnson, K. Kappel, J. Karanicolas, H. Khakzad, K. R. Khar, S. D. Khare, F. Khatib, A. Khramushin, I. C. King, R. Kleffner, B. Koepnick, T. Kortemme, G. Kuenze, B. Kuhlman, D. Kuroda, J. W. Labonte, J. K. Lai, G. Lapidoth, A. Leaver-Fay, S. Lindert, T. Linsky, N. London, J. H. Lubin, S. Lyskov, J. Maguire, L. Malmstrom, E. Marcos, O. Marcu, N. A. Marze, J. Meiler, R. Moretti, V. K. Mulligan, S. Nerli, C. Norn, S. O’Conchuir, N. Ollikainen, S. Ovchinnikov, M. S. Pacella, X. Pan, H. Park, R. E. Pavlovicz, M. Pethe, B. G. Pierce, K. B. Pilla, B. Raveh, P. D. Renfrew, S. S. R. Burman, A. Rubenstein, M. F. Sauer, A. Scheck, W. Schief, O. Schueler-Furman, Y. Sedan, A. M. Sevy, N. G. Sgourakis, L. Shi, J. B. Siegel, D. A. Silva, S. Smith, Y. Song, A. Stein, M. Szegedy, F. D. Teets, S. B. Thyme, R. Y. Wang, A. Watkins, L. Zimmerman, and R. Bonneau. 2020. Macromolecular modeling and design in Rosetta: recent methods and frameworks. Nat Methods. 17(7):665–680, doi: 10.1038/s41592-020-0848-2.

52. Schmitz, S., M. Ertelt, R. Merkl, and J. Meiler. 2021. Rosetta design with co-evolutionary information retains protein function. PLoS Comput Biol. 17(1):e1008568, doi: 10.1371/journal.pcbi.1008568.

53. Perkel, J. M. 2019. The computational protein designers. Nature. 571(7766):585–587, doi: 10.1038/d41586-019-02251-x.

54. Liu, Y., and B. Kuhlman. 2006. RosettaDesign server for protein design. Nucleic Acids Res. 34(Web Server issue):W235-238, doi: 10.1093/nar/gkl163.

55. Ertelt, M., V. K. Mulligan, J. B. Maguire, S. Lyskov, R. Moretti, T. Schiffner, J. Meiler, and C. T. Schoeder. 2024. Combining machine learning with structure-based protein design to predict and engineer post-translational modifications of proteins. PLoS Comput Biol. 20(3):e1011939, doi: 10.1371/journal.pcbi.1011939.

56. Fleishman, S. J., A. Leaver-Fay, J. E. Corn, E. M. Strauch, S. D. Khare, N. Koga, J. Ashworth, P. Murphy, F. Richter, G. Lemmon, J. Meiler, and D. Baker. 2011. RosettaScripts: a scripting language interface to the Rosetta macromolecular modeling suite. PLoS One. 6(6):e20161, doi: 10.1371/journal.pone.0020161.

57. Laio, A., and M. Parrinello. 2002. Escaping free-energy minima. Proceedings of the National Academy of Sciences of the United States of America. 99(20):12562–12566, doi: 10.1073/pnas.202427399.

58. Laio, A., and F. L. Gervasio. 2008. Metadynamics: a method to simulate rare events and reconstruct the free energy in biophysics, chemistry and material science. Reports on Progress in Physics. 71(12):126601, doi: 10.1088/0034-4885/71/12/126601.

59. Barducci, A., G. Bussi, and M. Parrinello. 2008. Well-tempered metadynamics: a smoothly converging and tunable free-energy method. Phys Rev Lett. 100(2):020603, doi: 10.1103/PhysRevLett.100.020603.

60. Dama, J. F., M. Parrinello, and G. A. Voth. 2014. Well-tempered metadynamics converges asymptotically. Phys Rev Lett. 112(24):240602, doi: 10.1103/PhysRevLett.112.240602.

61. Pearson, K. 1901. LIII. On lines and planes of closest fit to systems of points in space. The London, Edinburgh, and Dublin Philosophical Magazine and Journal of Science. 2(11):559–572, doi: 10.1080/14786440109462720.

62. Mendels, D., G. Piccini, and M. Parrinello. 2018. Collective Variables from Local Fluctuations. J Phys Chem Lett. 9(11):2776–2781, doi: 10.1021/acs.jpclett.8b00733.

63. Piccini, G., D. Mendels, and M. Parrinello. 2018. Metadynamics with Discriminants: A Tool for Understanding Chemistry. J Chem Theory Comput. 14(10):5040–5044, doi: 10.1021/acs.jctc.8b00634.

64. Tian, C., K. Kasavajhala, K. A. A. Belfon, L. Raguette, H. Huang, A. N. Migues, J. Bickel, Y. Wang, J. Pincay, Q. Wu, and C. Simmerling. 2020. h19SB: Amino-Acid-Specific Protein Backbone Parameters Trained against Quantum Mechanics Energy Surfaces in Solution. J Chem Theory Comput. 16(1):528–552, doi: 10.1021/acs.jctc.9b00591.

65. Izadi, S., R. Anandakrishnan, and A. V. Onufriev. 2014. Building Water Models: A Different Approach. J Phys Chem Lett. 5(21):3863–3871, doi: 10.1021/jz501780a.

66. Ryckaert, J.-P., G. Ciccotti, and H. J. C. Berendsen. 1977. Numerical integration of the cartesian equations of motion of a system with constraints: molecular dynamics of n-alkanes. Journal of Computational Physics. 23(3):327–341.

67. Forester, T. R., and W. Smith. 1998. SHAKE, rattle, and roll: Efficient constraint algorithms for linked rigid bodies. Journal of Computational Chemistry. 19(1):102–111, doi: Doi 10.1002/(Sici)1096-987x(19980115)19:1<102::Aid-Jcc9>3.0.Co;2-T.

68. Loncharich, R. J., B. R. Brooks, and R. W. Pastor. 1992. Langevin dynamics of peptides: the frictional dependence of isomerization rates of N-acetylalanyl-N’-methylamide. Biopolymers. 32(5):523–535, doi: 10.1002/bip.360320508.

69. Berendsen, H. J. C., J. P. M. Postma, W. F. Vangunsteren, A. Dinola, and J. R. Haak. 1984. Molecular-Dynamics with Coupling to an External Bath. Journal of Chemical Physics. 81(8):3684–3690, doi: Doi 10.1063/1.448118.

70. AMBER 2020 (University of California, San Francisco).

71. consortium,P. 2019. Promoting transparency and reproducibility in enhanced molecular simulations. Nat Methods. 16(8):670–673, doi: 10.1038/s41592-019-0506-8.

72. Tribello, G. A., M. Bonomi, D. Branduardi, C. Camilloni, and G. Bussi. 2014. PLUMED 2: New feathers for an old bird. Computer Physics Communications. 185(2):604–613, doi: 10.1016/j.cpc.2013.09.018.

73. Roe, D. R., and T. E. Cheatham, 3rd. 2013. PTRAJ and CPPTRAJ: Software for Processing and Analysis of Molecular Dynamics Trajectory Data. J Chem Theory Comput. 9(7):3084–3095, doi: 10.1021/ct400341p.

74. Humphrey, W., A. Dalke, and K. Schulten. 1996. VMD: visual molecular dynamics. J Mol Graph. 14(1):33-38, 27-38, doi: 10.1016/0263-7855(96)00018-5.

75. Park, H., P. Bradley, P. Greisen, Jr., Y. Liu, V. K. Mulligan, D. E. Kim, D. Baker, and F. DiMaio. 2016. Simultaneous Optimization of Biomolecular Energy Functions on Features from Small Molecules and Macromolecules. J Chem Theory Comput. 12(12):6201–6212, doi: 10.1021/acs.jctc.6b00819.

76. Efron, B. 1984. Better Bootstrap Confidence Intervals. J. Am. Stat. Assoc. 82(397):171–185, doi: 10.1080/01621459.1987.10478410.

77. Offer, J., C. N. Boddy, and P. E. Dawson. 2002. Extending synthetic access to proteins with a removable acyl transfer auxiliary. J Am Chem Soc. 124(17):4642–4646, doi: 10.1021/ja016731w.

78. Johnson, E. C. B., and S. B. H. Kent. 2006. Insights into the Mechanism and Catalysis of the Native Chemical Ligation Reaction. J Am Chem Soc. 128(20):6640–6646, doi: 10.1021/ja058344i.

79. Pace, J. R., B. J. Lampkin, C. Abakah, A. Moyer, J. Miao, K. Deprey, R. A. Cerulli, Y. S. Lin, J. D. Baleja, D. Baker, and J. A. Kritzer. 2021. Stapled beta-Hairpins Featuring 4-Mercaptoproline. J Am Chem Soc. 143(37):15039–15044, doi: 10.1021/jacs.1c04378.

80. Wang, Q., F. Wang, R. Li, P. Wang, R. Yuan, D. Liu, Y. Liu, Y. Luan, C. Wang, and S. Dong. 2023. Fine Tuning the Properties of Stapled Peptides by Stereogenic alpha-Amino Acid Bridges. Chemistry. 29(29):e202203624, doi: 10.1002/chem.202203624.

81. Bhardwaj, G., V. K. Mulligan, C. D. Bahl, J. M. Gilmore, P. J. Harvey, O. Cheneval, G. W. Buchko, S. V. Pulavarti, Q. Kaas, A. Eletsky, P. S. Huang, W. A. Johnsen, P. J. Greisen, G. J. Rocklin, Y. Song, T. W. Linsky, A. Watkins, S. A. Rettie, X. Xu, L. P. Carter, R. Bonneau, J. M. Olson, E. Coutsias, C. E. Correnti, T. Szyperski, D. J. Craik, and D. Baker. 2016. Accurate de novo design of hyperstable constrained peptides. Nature. 538(7625):329–335, doi: 10.1038/nature19791.

82. Schroder, H. V., M. Stadlmeier, M. Wuhr, and A. J. Link. 2022. The Shuttling Cascade in Lasso Peptide Benenodin-1 is Controlled by Non-Covalent Interactions. Chemistry. 28(5):e202103615, doi: 10.1002/chem.202103615.

83. Al Toma, R. S., A. Kuthning, M. P. Exner, A. Denisiuk, J. Ziegler, N. Budisa, and R. D. Sussmuth. 2015. Site-directed and global incorporation of orthogonal and isostructural noncanonical amino acids into the ribosomal lasso peptide capistruin. Chembiochem. 16(3):503–509, doi: 10.1002/cbic.201402558.

