## Supplementary Material for "Tying the Knot: In Silico Design of Foldable Lasso Peptides"

**Listing S1.** Sample RosettaScripts script used for fixed-backbone sequence design.

---

```
<ROSETTASCRIPTS>
 #scoring functions that will be used later in the script
    <SCOREFXNS>
      <ScoreFunction name="ref" weights="ref2015" />
      <ScoreFunction name="ref_cst" weights="ref2015" >
     <Reweight scoretype="atom_pair_constraint" weight="1.0" />
     <Reweight scoretype="dihedral_constraint" weight="1.0" />
     <Reweight scoretype="angle_constraint" weight="1.0" />
     </ScoreFunction>
    </SCOREFXNS>
#custom palette to sample D-AAs
    <PACKER_PALETTES>
      <CustomBaseTypePackerPalette name="custom_palette"
additional_residue_types="DALA,DASP,DGLU,DPHE,DHIS,DILE,DLYS,DLEU,DMET
,DASN,DPRO,DGLN,DARG,DSER,DTHR,DVAL,DTRP,DTYR" />
    </PACKER_PALETTES>
    <TASKOPERATIONS>
      <ReadResfile name="resfile" filename="resfile.txt" />
    </TASKOPERATIONS>
#filter to discard designs with more than two hydrogen bonds to
carbonyls (score function artifact)
    <FILTERS>
      <OversaturatedHbondAcceptorFilter name="oversat" scorefxn="ref"
max_allowed_oversaturated="0" consider_mainchain_only="false" />
    </FILTERS>
#isopeptide bond torsion, angle, and distance constraints
    <MOVERS>
      <DeclareBond name="connect_1_to_8" res1="1" res2="8" atom1="N"
atom2="CD" />
      <CreateTorsionConstraint name="peptide_torsion_constraint">
     <Add res1="1" res2="1" res3="8" res4="8" atom1="CA" atom2="N"
atom3="CD" atom4="CG" cst_func="CIRCULARHARMONIC 3.1360076 0.005" />
     <Add res1="1" res2="1" res3="8" res4="8" atom1="CA" atom2="N"
atom3="CD" atom4="OE" cst_func="CIRCULARHARMONIC -0.008203047 0.005"
/>
```

```
      <Add res1="1" res2="1" res3="8" res4="8" atom1="H" atom2="N"
atom3="CD" atom4="OE" cst_func="CIRCULARHARMONIC -3.14071999 0.005" />
      <Add res1="1" res2="1" res3="8" res4="8" atom1="H" atom2="N"
atom3="CD" atom4="CG" cst_func="CIRCULARHARMONIC 0.00349066 0.005" />
      </CreateTorsionConstraint>
      <CreateAngleConstraint name="peptide_angle_constraints">
      <Add res1="1" atom1="CA" res_center="1" atom_center="N" res2="8"
atom2="CD" cst_func="CIRCULARHARMONIC 2.11219746 0.005" />
      <Add res1="1" atom1="N" res_center="8" atom_center="CD" res2="8"
atom2="CG" cst_func="CIRCULARHARMONIC 2.04308242 0.005" />
      <Add res1="1" atom1="N" res_center="8" atom_center="CD" res2="8"
atom2="OE" cst_func="CIRCULARHARMONIC 2.16193934 0.005" />
      <Add res1="1" atom1="H" res_center="1" atom_center="N" res2="8"
atom2="CD" cst_func="CIRCULARHARMONIC 2.08095607 0.005" />
      </CreateAngleConstraint>
      <CreateDistanceConstraint name="1_to_8_dist_cst">
      <Add res1="1" res2="8" atom1="N" atom2="CD" cst_func="HARMONIC
1.331084 0.01" />
      </CreateDistanceConstraint>
#flexible backbone design
      <PackRotamersMover name="prm" scorefxn="ref_cst" nloop="3"
task_operations="resfile" packer_palette="custom_palette" >
      </PackRotamersMover>
    </MOVERS>
    <PROTOCOLS>
      <Add mover="connect_1_to_8" />
      <Add mover="peptide_torsion_constraint" />
      <Add mover="peptide_angle_constraints" />
      <Add mover="1_to_8_dist_cst" />
      <Add mover="prm" />
         <Add mover="connect_1_to_8" />
       <Add mover="connect_1_to_8" />
         <Add filter="oversat" />
    </PROTOCOLS>
    <OUTPUT scorefxn="ref" />
</ROSETTASCRIPTS>
```

**Listing S2.** Sample PLUMED input file used for WTMetaD.

```
# native MJ25
RESTART

# units
UNITS LENGTH=A

# Molecule
WHOLEMOLECULES ENTITY0=1-292
```

```
# c-aplha distances
d1: DISTANCE ATOMS=5,246
d2: DISTANCE ATOMS=12,246
d3: DISTANCE ATOMS=19,246
d4: DISTANCE ATOMS=29,246
d5: DISTANCE ATOMS=36,246
d6: DISTANCE ATOMS=53,246
d7: DISTANCE ATOMS=77,246
d8: DISTANCE ATOMS=83,246

# coordination numbers
1_8: COORDINATION GROUPA=1 GROUPB=91 R_0=6.0 NOPBC
6_20: COORDINATION GROUPA=66 GROUPB=264 R_0=4.0 NOPBC

CV1: COMBINE ...
        ARG=d1,d2,d3,d4,d5,d6,d7,d8
        COEFFICIENTS=-0.5128,-0.2235,-0.1513,-0.2441,-0.1942,-0.3969,-
0.5767,-0.277
        PERIODIC=NO
...

#1_8: DISTANCE ATOMS=1,91
#6_20: DISTANCE ATOMS=66,264


CV2: COMBINE ...
        ARG=1_8,6_20
        PERIODIC=NO
...

# well-tempered metadynamics
metad: METAD ARG=CV1,CV2 ...
   PACE=500 HEIGHT=0.03 BIASFACTOR=10 TEMP=300
   # Gaussian width (sigma) should be chosen based on the CV
fluctuations in unbiased run
   # 1/2 of standard deviation of CV is used here
   SIGMA=0.11,0.075
   # Gaussians will be written to file and also stored on grid
   FILE=HILLS
   GRID_WSTRIDE=100000
GRID_RFILE=grid01.dat
GRID_WFILE=grid02.dat
   GRID_NOSPLINE
   GRID_MIN=-120,0
   GRID_MAX=0,2
   GRID_BIN=1200,20
...
# Print both collective variables on COLVAR file every 100 steps
PRINT ARG=CV1,CV2,metad.work,metad.bias FILE=COLVAR STRIDE=100
```

---

**Table S1.** Rosetta-designed sequences designed from PDB structures of class II lasso peptides. Lowercase letters in a sequence represent D-amino acids. REF2015 was used to score the lasso structure of each sequence.

**Rosetta-designed sequences**

| Name | PDB ID | Native sequence<br><br>Designed sequence | Sequence length | Ring length | Loop length | Tail length | Score (REU) | Δ Native (REU) |
|---|---|---|---|---|---|---|---|---|
| rubrivinodin | 5OQZ | GAPSLINSEDNPAFPQRV<br><br>GRPDSDNSENNPDHPAAQ | 18 | 9 | 7 | 2 | -37.00 | -9.18 |
| caulosegnin II | 5D9E | GAFVGQPEAVNPLGREIQG<br><br>GDSQPaLPEETKPGHFIPS | 19 | 9 | 6 | 4 | -36.39 | -4.21 |
| achromonodin-1 | 8SVB | GGGGPTPEYFLMPIDPAWLQANLPNTGKYN<br><br>GtGGPGPENELNPDDPDFRNDNKPQTGYYS | 30 | 8 | 20 | 2 | -31.69 | -19.50 |
| microcin J25 | 1Q71 | GGAGHVPEYFVGIGTPISFYG<br><br>GsHsKIAEEFDKDGVPRSWYG | 21 | 8 | 11 (β-hairpin) | 2 | -31.17 | -6.01 |
| caulonodin V | 2MLJ | SIGDSGLRESMSSQTYWP<br><br>SNGDSSVPESSESNKYYE | 18 | 9 | 7 | 2 | -28.29 | -8.50 |
| caulosegnin I | 2LX6 | GAFVGQPEAVNPLGREIQG<br><br>GDSEGKPETTEPKGHVKES | 19 | 8 | 7 | 4 | -22.91 | -12.21 |
| streptomonomicin | 2MW3 | SLGSSPYNDILGYPALIVIYP<br><br>SDGSAPETDEKGKPSNVVNYP | 21 | 9 | 5 | 7 | -16.79 | -14.04 |
| astexin-1 | 2LTI | SLGSSPYNDILGYPALIVIYP<br><br>GGNVGTEPDDGQTDKHYDRRNQS | 23 | 9 | 8 | 6 | -14.70 | -28.44 |
| acinetodin | 5UI6 | GGKGPIFETWVTEGNYYG<br><br>GGSyDDPEHEVKNGTLYG | 18 | 8 | 8 | 2 | -9.68 | -14.94 |
| chaxapeptin | 2N5C | GFGSKPLDSFGLNFF<br><br>GEGDQPSDDKsLASQ | 15 | 8 | 4 | 3 | -9.26 | -8.50 |
| citrocin | 6MW6 | GGVGKIIEYFIGGGVGRYG<br><br>GGEGKEDEASHsGGKGKFa | 19 | 8 | 9 | 2 | -2.55 | -8.19 |

| | | | | | | | | |
|---|---|---|---|---|---|---|---|---|
| lihuanodin | 7LCW | GSKYSDTADESSYRW<br><br>GSGPSGTDDDNAASS | 15 | 9 | 4 | 2 | -2.46 | -4.07 |
| astexin-3 | 2M8F | GPTPMVGLDSVSGQYW<br>DQHAPLAD<br><br>GSKPMDGGDEETGGKW<br>RGSAnKAE | 24 | 9 | 6 | 9 | -2.06 | -0.97 |
| stlassin | 7BZA | LVVIVQADWNAPGWF<br><br>LTHRGGSDEKRPGFE | 15 | 8 | 5 | 2 | -1.59 | -15.52 |
| triculamin | 7ZWJ | SKKSKPGDGIRGKGVR<br>G<br><br>SHSSAPEDGDGGAGKK<br>s | 17 | 8 | 6 | 3 | 2.98 | -23.16 |
| rubrinodin | 7EES | GTIDPQNSEEHPVLSR<br>RLEN<br><br>GKVEPKNTEPEPDEPA<br>AQYE | 20 | 9 | 7 | 4 | 9.76 | -12.32 |
| des-citrulassin F | 7JS6 | LLGRSGNDRLILSKN<br><br>LNGNSGNDdGTASEE | 15 | 8 | 3 | 4 | 16.73 | -10.10 |

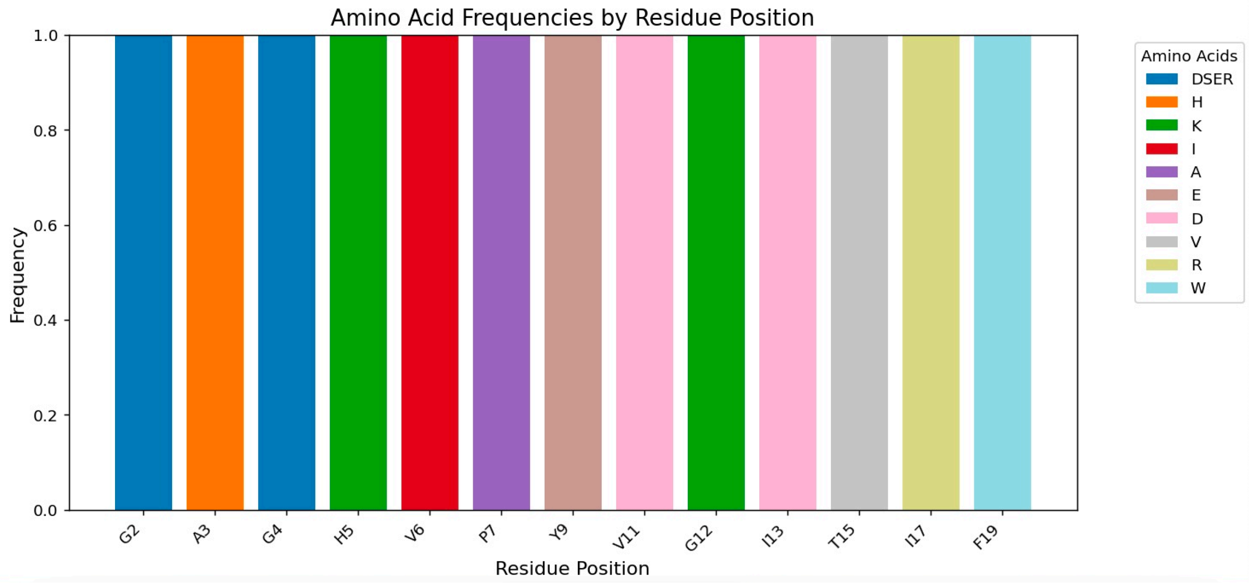


**Figure S1.** Sequence changes in MccJ25 from design with the NMR solution structure.

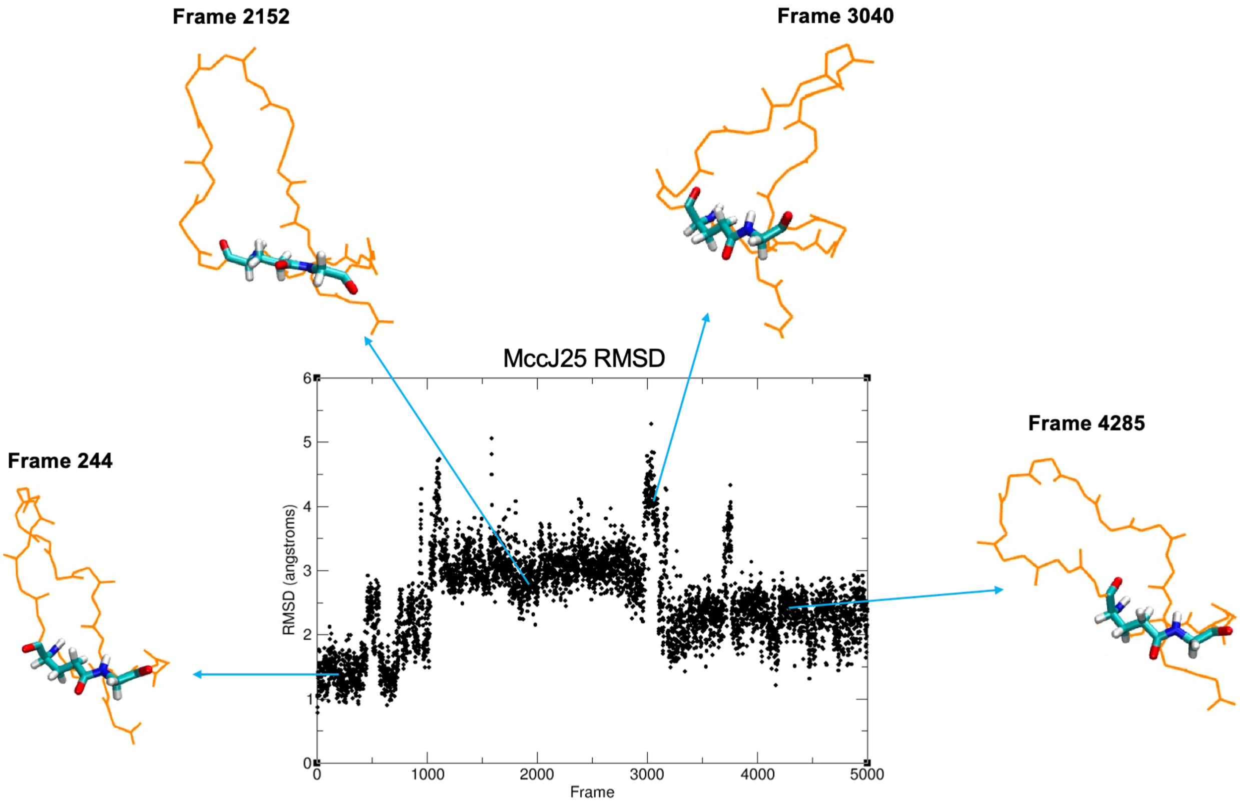


**Figure S2.** Conformations of native McCJ25 throughout a 5000 ns standard MD simulation that were used for additional fixed-backbone design runs.

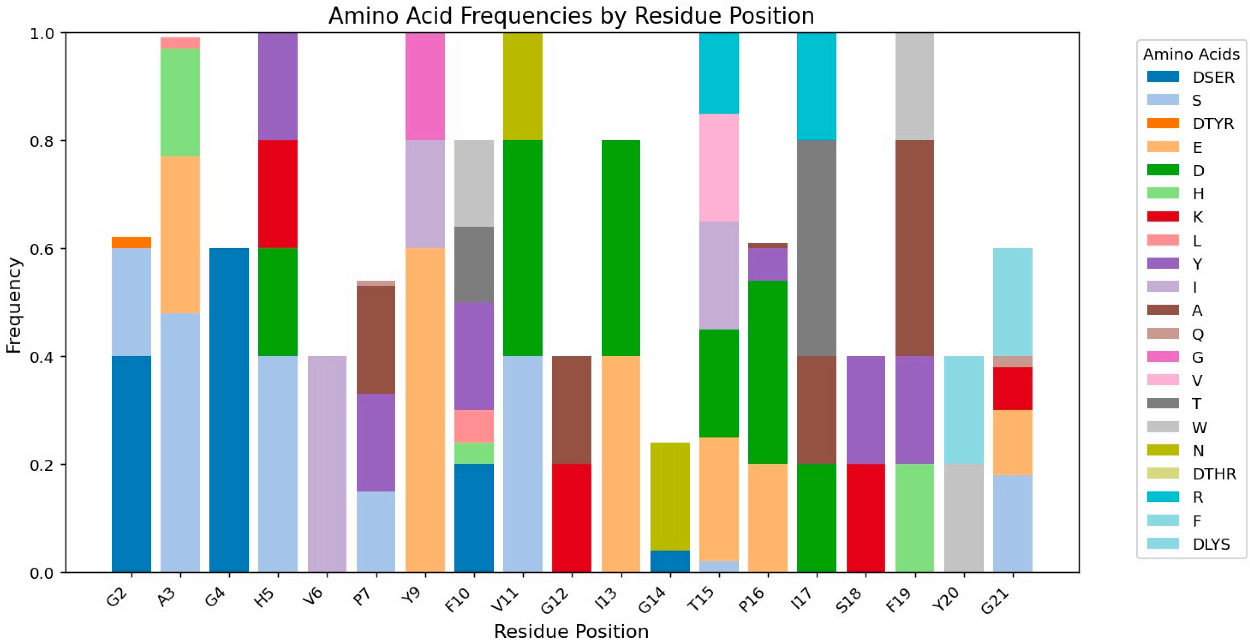


**Figure S3.** Sequence changes in MccJ25 from design with the NMR solution structure along with the four additional MD structures in **Figure S2**.

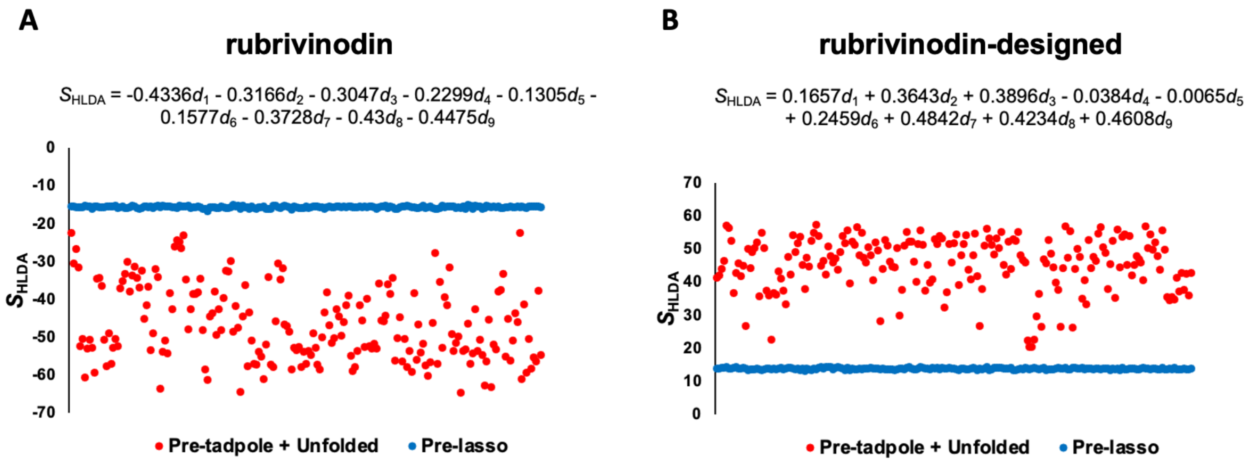


**Figure S4.** HLDA analysis for native (A) and designed (B) sequences of Rubrivinodin. The linear combination of C-alpha distances for each sequence is shown.

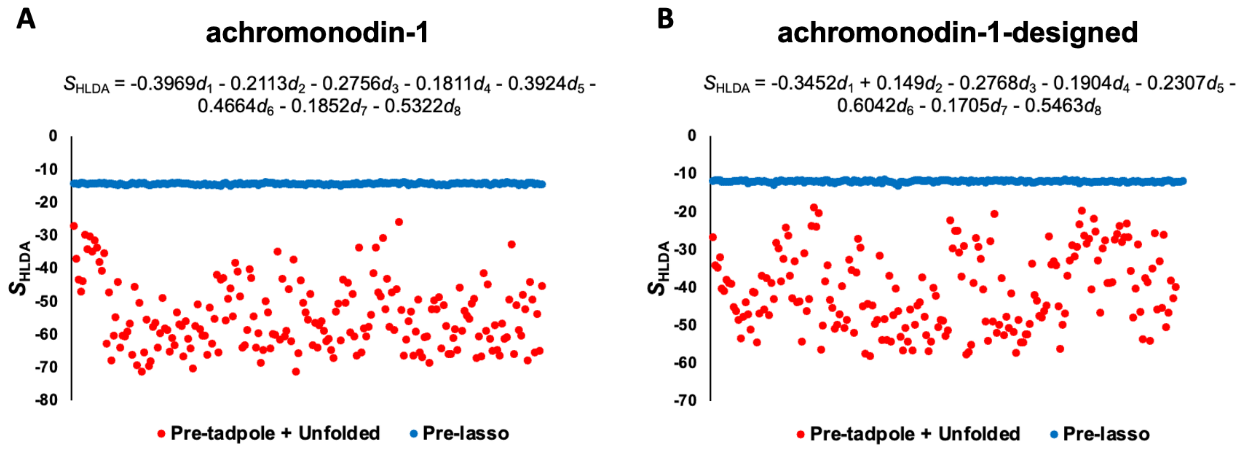


**Figure S5.** HLDA analysis for native (A) and designed (B) sequences of Achromonodin-1. The linear combination of C-alpha distances for each sequence is shown.

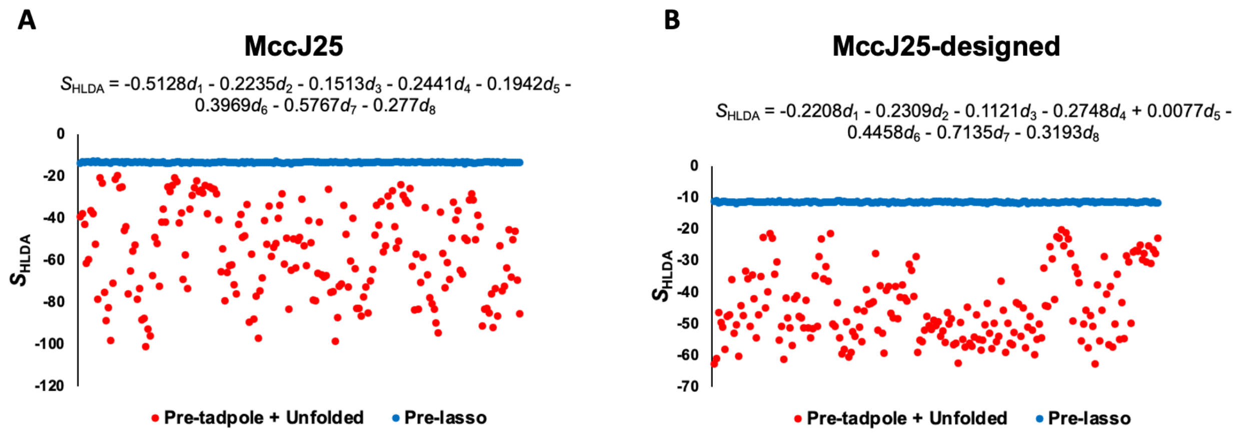

**Figure S6.** HLDA analysis for native (A) and designed (B) sequences of Microcin J25. The linear combination of C-alpha distances for each sequence is shown.

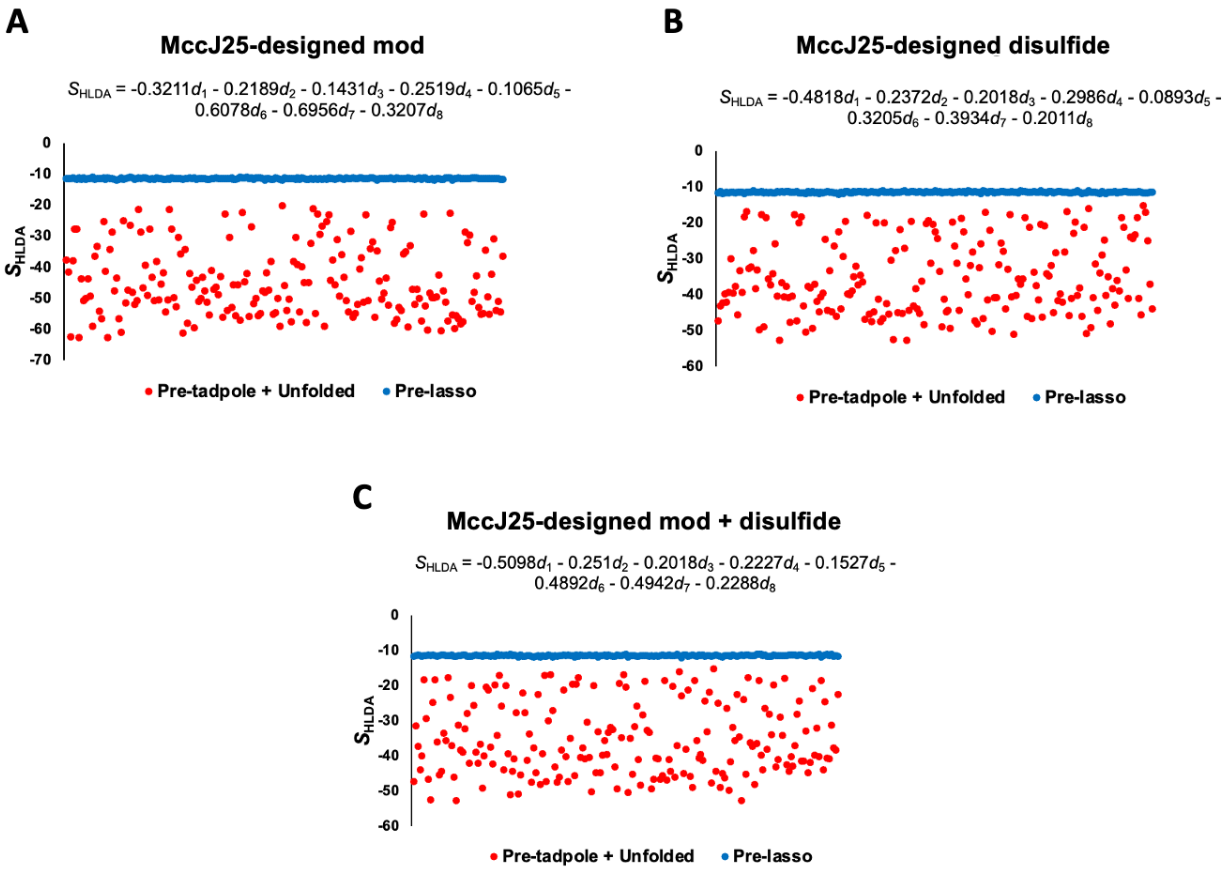


**Figure S7.** HLDA analysis for designed + modifications (A), designed + disulfide (B), and designed + modifications + disulfide (C) sequences of Microcin J25. The linear combination of C-alpha distances for each sequence is shown.

**Table S2.** Standard errors for the pre-lasso and pre-tadpole probabilities from **Tables 1** and **2**. Error is calculated by bias-corrected bootstrapping. Designed sequences are represented with (-d).

| Name | Pre-lasso probability (%) | Pre-tadpole probability (%) |
|---|---|---|
| Rubrivinodin | $9.18 \times 10^{-3} \pm 6.55 \times 10^{-4}$ | $0.83 \pm 0.02$ |
| Rubrivinodin-d | $0.13 \pm 0.01$ | $0.45 \pm 6.98 \times 10^{-4}$ |
| Achromonodin-1 | $2.54 \times 10^{-6} \pm 2.86 \times 10^{-8}$ | $9.89 \times 10^{-6} \pm 4.02 \times 10^{-9}$ |
| Achromonodin-1-d | $5.07 \times 10^{-5} \pm 1.06 \times 10^{-6}$ | $9.15 \times 10^{-4} \pm 1.17 \times 10^{-5}$ |

| | | |
|---|---|---|
| MccJ25 | $3.63 \times 10^{-4} \pm 5.25 \times 10^{-6}$ | 0.94 ± 0.01 |
| MccJ25-d | $1.31 \times 10^{-2} \pm 1.22 \times 10^{-3}$ | 0.70 ± 0.01 |
| MccJ25-d mod | 1.40 ± 0.3 | 8.53 ± 0.6 |
| MccJ25-ddisulfide | $9.56 \times 10^{-4} \pm 2.72 \times 10^{-5}$ | 2.01 ± 0.06 |
| MccJ25-d mod + disulfide | 3.26 ± 0.6 | 12.57 ± 0.9 |

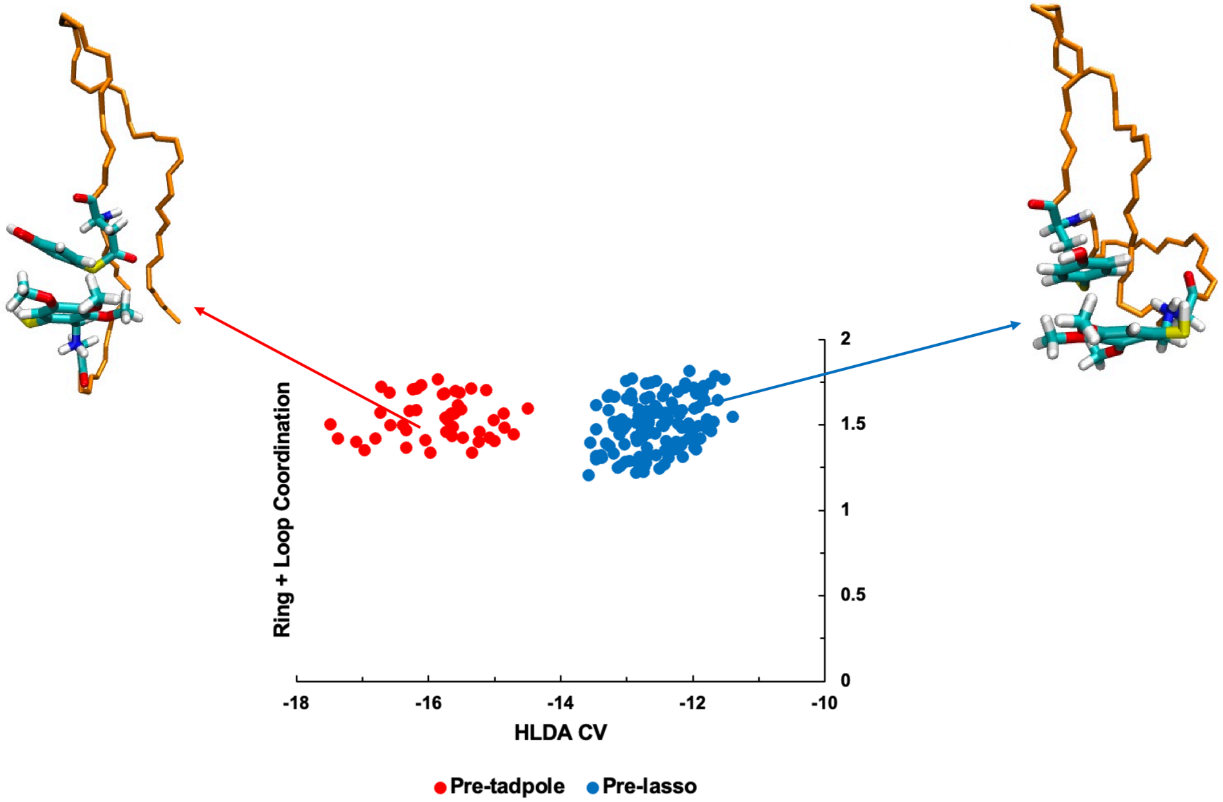


**Figure S8.** The distribution of pre-lasso and pre-tadpole structures in **Figure 9B** focusing on an HLDA CV range of -18 to -10 to show the separation between the pre-lasso and pre-tadpole that closely resembles the pre-lasso.
